# Dissecting Immune–Epithelial Interactions in Airway Infection at Single-Cell Resolution Using a Compartmentalised Microfluidic Device

**DOI:** 10.64898/2026.08.24.746128

**Authors:** Lucy-May G. Young, Christopher P. Tostado, Jie-Yi Koh Kok, Jorge Amaya Cataño, Ramanuj DasGupta, Kirsten M. Spann, Yi-Chin Toh

**Author notes:** Corresponding author(s) **-** Y.-C. Toh, - K.M. Spann.

## Abstract

Immune–epithelial interactions govern the initiation and progression of airway diseases, yet their heterogeneity is difficult to capture using existing in vitro models. Although conventional Transwell^®^ and lung-on-chip systems reproduce airway compartmentalisation and permit epithelial-immune interactions, they lack the spatial and analytical resolution needed to visualise dynamic immune behaviour during infection. Here, we present the “Single Cell resolved Airway-Immune Recruitment” (scAIR) platform designed to interrogate immune–epithelial interactions during airway infection. The scAIR device features a modular central chamber accommodating a Transwell^®^ insert with primary airway epithelial cells (AECs) pre-differentiated under air–liquid interface (ALI), flanked by immune compartments connected through a precision-patterned microchannel array. This architecture enables real-time single-cell imaging of immune cell migration while preserving epithelial physiology. The scAIR device coupled with a machine learning analysis (MLA) pipeline enables automated tracking and quantification of individual immune cell speed, direction, and behavioural heterogeneity. Using this platform, respiratory syncytial virus (RSV) infection is modelled to generate a type 1 inflammatory airway epithelium that drives neutrophil recruitment. TNF-α neutralisation with adalimumab reveals distinct migratory behaviours that are obscured by population-averaged measurements. This integrated platform quantifies airway immune responses during infection and therapeutic modulation, enabling mechanistic studies, drug evaluation, and precision modelling of airway inflammation.

## 1. Introduction

The human respiratory airways provide a direct interface with the inhaled environment, serving as a first line of defence against incoming pathogens and particulates. Besides serving as a physical barrier to environmental insults, the airway epithelium also plays a crucial role in orchestrating the resultant immune responses that drive the progression and resolution of disease. ^[1]^ However, immune responses within the respiratory airways are highly heterogeneous, varying across disease contexts and are shaped by host-specific factors such as age and co-morbidities. In chronic airway diseases, including asthma, chronic obstructive pulmonary disease (COPD), and chronic rhinosinusitis (CRS), this heterogeneity is reflected in distinct inflammatory endotypes, most commonly characterised as type 2 (eosinophilic) ^[2–4]^ or type 1 (neutrophilic) ^[5–7]^ inflammation, which arise from fundamentally different immune pathways during exacerbation. Similar levels of non-uniformity are observed during acute viral respiratory infections such as respiratory syncytial virus (RSV) and influenza, which predominantly induce a type 1 immune response in healthy adults, but elicit a preference for a type 2 response in infants and the elderly. ^[8, 9]^ Such immune heterogeneity presents a major challenge for developing *in vitro* models capable of predicting disease severity and therapeutic responsiveness. Effective airway models must not only provide a physiologically relevant microenvironment that recapitulates the spatial-temporal dynamics of pathogen entry, epithelial sensing, and immune activation, but also enable resolution of variability in immune behaviour across diverse inflammatory contexts.

To date, *in vitro* airway models have primarily focused on replicating the physiological microenvironment of the airway epithelium, including cellular composition and sources, as well as the mucosal architecture, rather than on resolving immune heterogeneity. The prevailing approach involves coculturing airway epithelial cells (AECs) at the air–liquid interface (ALI) on a semi-permeable membrane, with endothelial cells and/or immune cells seeded in the basal compartment. This physical compartmentalisation of cell populations is typically achieved using traditional static Transwell^®^ systems ^[10, 11]^ or microfluidic lung-on-chip (LOC) platforms that adopt a similar top-bottom configuration but with a dynamic culture setup. ^[12–14]^ Both static and dynamic platforms have been shown to support long-term differentiation (typically > 21 days) of AECs into well-polarised epithelium with intact barrier and mucociliary functions, which are suitable for modelling pathogen infection and epithelial responses. ^[12, 15]^ However, these systems are intrinsically limited for studying immune dynamics because the vertically stacked apical–basal configuration and the absence of defined migration paths restrict measurements to bulk, endpoint readouts, obscuring real-time quantitative analysis of the sequential and heterogeneous nature of immune recruitment to the airway epithelium.

Another class of microfluidic airway models features a horizontally-orientated apical–basal configuration aligned in a single imaging plane, which facilitates real-time visualisation of pathogen invasion and cell migration across the airway epithelium. Compartmentalisation is typically achieved by patterning a hydrogel matrix (e.g., collagen) using microstructures such as phase guides or micropillars. ^[16–20]^ AECs are seeded along the hydrogel surface, while stromal cells, such as pulmonary fibroblasts, and vascular cells, can be embedded within the matrix. While these systems have been used to study airway epithelium responses, including inflammatory cytokine and mucous production, epithelium and endothelium remodelling, as well as leukocyte recruitment to viral, bacterial, and fungal infections ^[16, 21]^, they remain underutilised for dissecting heterogeneity in immune responses to epithelial-derived signals during infection. This gap is largely due to the technical challenges in synchronising AEC differentiation, pathogen exposure, immune cell loading, and high-resolution single-cell tracking within a single integrated system, which often limit reproducibility, scalability, and broader adoption for mechanistic and translational studies.

To overcome these limitations and enable the study of immune heterogeneity, an *in vitro* airway model must integrate robust epithelial differentiation with real-time, single-cell analysis of immune behaviour across dynamic inflammatory states. ^[22, 23]^ Here, we introduce the Single Cell resolved Airway-Immune Recruitment (termed here “scAIR”) platform engineered to meet this need. The platform comprises a compartmentalised microfluidic device with a horizontally aligned architecture, in which two immune compartments flank a central airway chamber, separated by a precision-patterned microchannel network. Integration of commercial Transwell^®^ inserts supports long-term ALI-dependent differentiation of AECs while decoupling epithelium maturation from downstream immune interaction assays, thereby improving reproducibility and experimental control. The microchannel network provides a geometrically defined migration zone for real-time, single-cell imaging of immune recruitment, offering greater precision for quantifying cell migration than hydrogel-based barriers. ^[24–26]^ A machine learning analysis (MLA) pipeline is implemented for automated, high-throughput single-cell tracking, enabling quantitative analysis of immune cell velocity, direction, and behavioural heterogeneity. Using this integrated system, we modelled RSV-induced type 1 airway inflammation, neutrophil recruitment and TNF-α signialling neutralisation with adalimumab, revealing subtle yet distinct immune behaviours that remain undetectable in conventional bulk assays during these processes. Together, these advances establish our platform as a robust and scalable tool for dissecting the complexity of airway–immune interactions and for uncovering how infections and immunomodulatory therapies reshape immune dynamics at single-cell resolution.

## 2. Results

The scAIR device was designed to model the spatial and temporal interaction between airway epithelial cells and immune cells under inflammatory conditions. By physically separating a functional airway epithelium from suspended immune cells, the sequential process of pathogen entry, epithelial-driven sensing, and subsequent immune cell recruitment to the infected airway epithelium can be realised, mimicking the tightly regulated process observed *in vivo* **(Figure 1A**).

**Figure 1.**
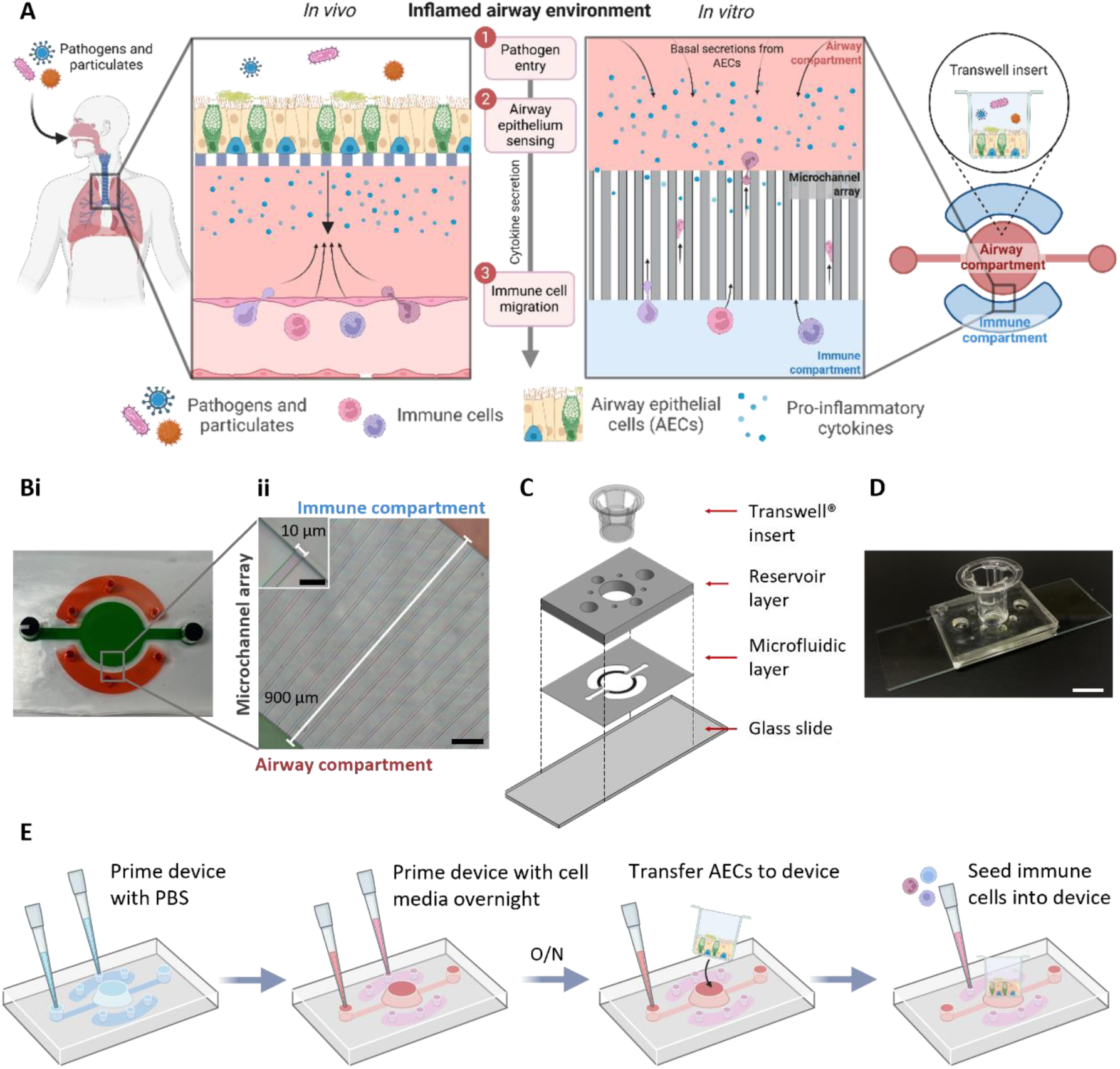
Development of a compartmentalised airway-immune interaction model. (A) Schematic representation of the inflamed airway environment and its recreation in this study using the scAIR device. (B) (i) Representative image of the microfluidic layer of the scAIR device patterned with coloured dyes, showing the immune (red) and airway (green) compartments. (ii) Magnified view of the microchannel network connecting the two compartments. Scale bar represents 100 µm. (C) Exploded view of the multilayered scAIR device. (D) Representative image of the fabricated device with Transwell^®^ insert. Scale bar represents 10 mm. (E) Schematic showing the process of performing airway-immune interaction assay using the scAIR device.

The scAIR device comprised a central airway compartment (green) separated from two flanking immune compartments (red) (**Figure 1B (i)**), which can host airway epithelial cells and immune cells, respectively. Although these compartments appear physically separated from one another, they are linked by an array of microchannels that enable the diffusion of soluble factors and the migration of immune cells. We designed the microchannels to be 900 × 10 x 5 µm (*l x w x h*) to ensure the cross-sectional size was small enough to restrict random immune cell passage but be permissive to chemokine-driven immune cell migration (**Figure 1B (ii))**.

AECs are predominantly cultured on suspended semi-permeable membranes (e.g., Transwell^®^ inserts) at an air-liquid interface (ALI), which allows access to both apical and basolateral compartments. Instead of integrating a precisely cut semi-permeable membrane into the microfluidic device, which would require the AECs to be differentiated on-device for >3 weeks, we configured the scAIR device to be compatible with commercially available Transwell^®^ inserts. This was achieved by adding a reservoir layer to sit atop the functional microfluidic layer and provide stable housing for the Transwell^®^ insert (**Figure 1C-D)**. This open-well design enabled us to pre-differentiate AECs off-device for >3 weeks before inserting the cell-laden Transwell^®^ insert into the device for downstream airway-immune interaction assays (**Figure 1E)**.

### 2.1. The scAIR device supports a well-differentiated AEC culture at the air–liquid interface

AECs are routinely cultured at the air-liquid interface (ALI) to facilitate mature differentiation into a pseudostratified epithelium containing ciliated, goblet and basal cells, which are tightly knitted together by tight junction complexes. Cells at ALI receive nutrients solely from culture medium at the basolateral side of the semi-permeable membrane. In traditional well-plate configurations, ALI cultures are typically maintained with 600 µL of basal media. Here, we designed the scAIR device to support AEC culture with reduced volume (i.e., ∼100 µL) of culture medium in the basolateral compartment with the goal of generating more concentrated secreted cytokines for downstream recruitment assays (**Figure 2A)**. To confirm the reduced cell medium volume could maintain a mature, differentiated airway epithelium, we transferred pre-differentiated AECs into scAIR devices and continued ALI culture for a total of 5 days. We found that cells retained high viability in scAIR devices when compared to well-plate culture, despite the lower volume of medium (**Figure 2B).** However, device cultures required more frequent media replenishment than conventional well-plate cultures, with media exchange performed every 24 hours rather than every 48 hours, respectively. Next, we assessed how well epithelial barrier integrity was maintained in the scAIR devices by measuring trans-epithelial electrical resistance (TEER) pre-and post-device culture. We found that the integrity of the monolayer remained consistent after 5 days of device culture compared with that at day 0 and noted no significant difference in TEER values between AECs cultured in scAIR devices, and those in well-plate cultures (**Figure 2C)**. This result also suggests that transfer of pre-differentiated AECs into devices was non-disruptive to the integrity of the cell monolayer.

**Figure 2.**
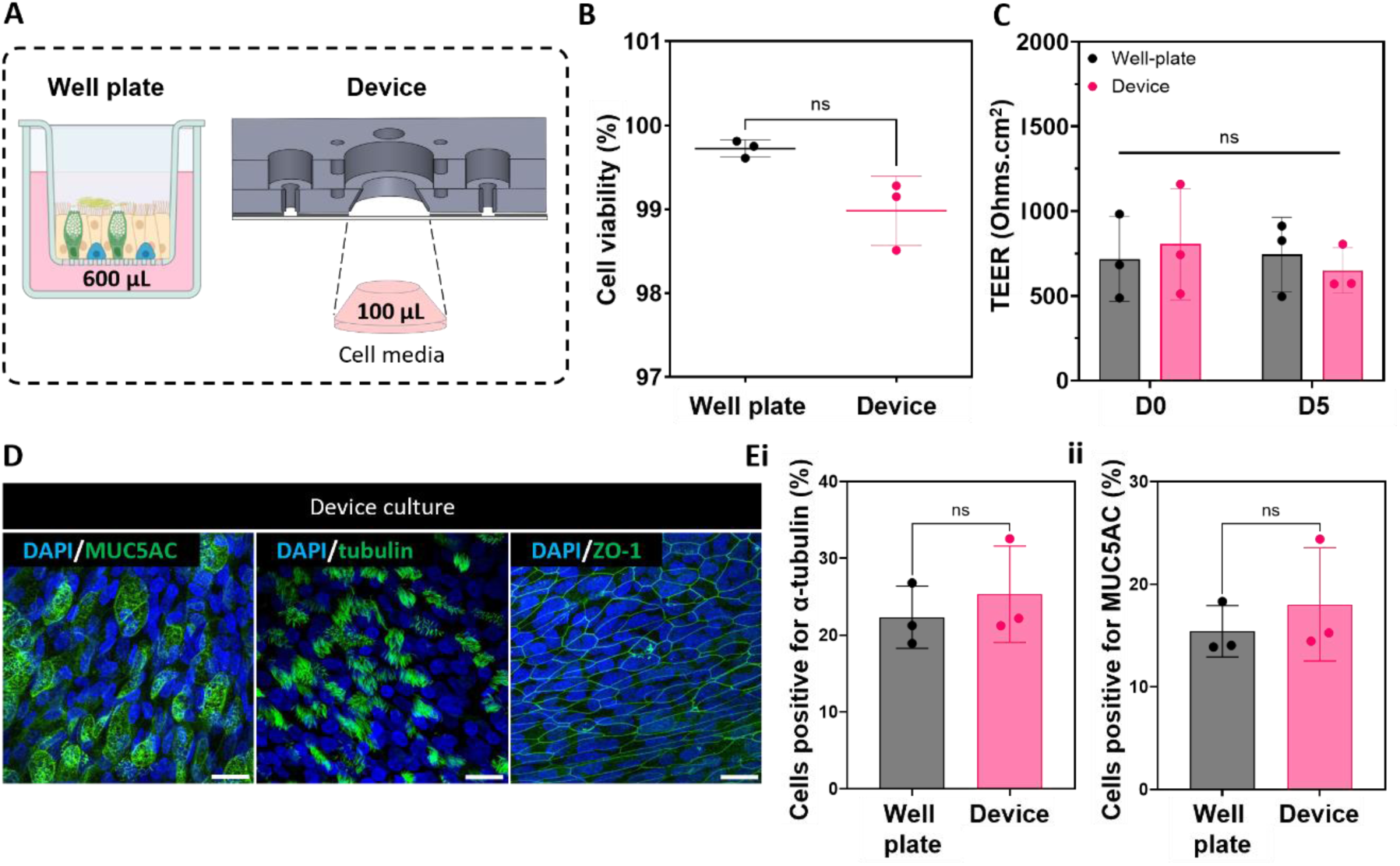
The scAIR device maintains differentiated airway epithelial phenotype and barrier integrity. (A) Schematic comparison of basal media volumes provided to AECs cultured on traditional Transwell® inserts and in scAIR devices. (B) Quantification of cell viability following live/dead staining. Data are presented as mean ± s.d.; n = 3 cultures per condition. Statistical significance was determined using an unpaired t-test with Welch’s correction. ns: not significant. (C) Barrier integrity of AECs following 5 days of culture in the scAIR device or well plates. Data are presented as mean ± s.d.; n = 3 cultures per condition. Statistical significance was determined using a two-way ANOVA with multiple comparisons. ns: not significant. (D) Representative immunofluorescence images of goblet cells (MUC5AC), ciliated cells (α-tubulin), and tight junctions (ZO-1) in AECs following 5 days of device culture. Scale bar, 100 µm. (E) Quantification of cells positive for α-tubulin (i) and MUC5AC (ii) staining in device culture compared with well-plate culture. n = 3 cultures per condition. Statistical significance was determined using an unpaired t-test with Welch’s correction. ns: not significant.

An advantage of the modular design of the scAIR device was that the entire Transwell^®^ insert could be removed post-experiment to facilitate confocal imaging without facing technical limitations such as working distance and optical distortion often observed in traditional closed-well microfluidic systems. We focused on measuring key functional markers of a mature, differentiated AEC monolayer, namely mucin5AC (MUC5AC), alpha-acetylated tubulin (α-acetylated tubulin), and zonula occludens-1 (ZO-1), which would indicate the presence of goblet cells, ciliated cells and tight junctions, respectively (**Figure 2D)**. We observed that expression levels of these key differentiation markers were comparable between AECs cultured in devices and well-plate control, thereby providing us with confidence that, despite the reduced basolateral volume of cell media, the scAIR supported the maintenance of a mature, differentiated airway epithelium for extended culture periods (**Figure 2E (i-ii)).**

### 2.2. A precision-engineered microchannel barrier enables chemotactic gradient formation and single-cell immune tracking

To enable communication between the airway and immune compartments in scAIR devices, we specifically designed the dimensions of the microchannel barrier to permit immune cells to transit only through active cell migration. It was also important to ensure that the microchannel geometry prevented convective mixing, thereby enabling the establishment of concentration gradients of soluble factors, such as cytokines and chemokines, between the airway and immune compartments to drive immune cell recruitment. To evaluate this, we measured the diffusion kinetics of 10 and 20 kDa FITC–dextran molecules, which approximate the molecular weights of common chemotactic factors ^[27]^, from the airway compartment to the immune compartment via the microchannel array over 24 hours (**Figure 3A**). We found that both 10 and 20 kDa FITC-Dextran were detected in the immune compartments at comparable rates over the first 6 hours. As expected, the smaller 10 kDa FITC-dextran exhibited a significantly higher rate of diffusion across the microchannel array than 20 kDa FITC-dextran from 6-24 hours (**Figure 3B**), consistent with molecular size-dependent passive diffusion. These results showed that soluble factor gradients could be established across the microchannels in scAIR devices within timeframes relevant to early immune signalling during infection. ^[28–30]^

**Figure 3.**
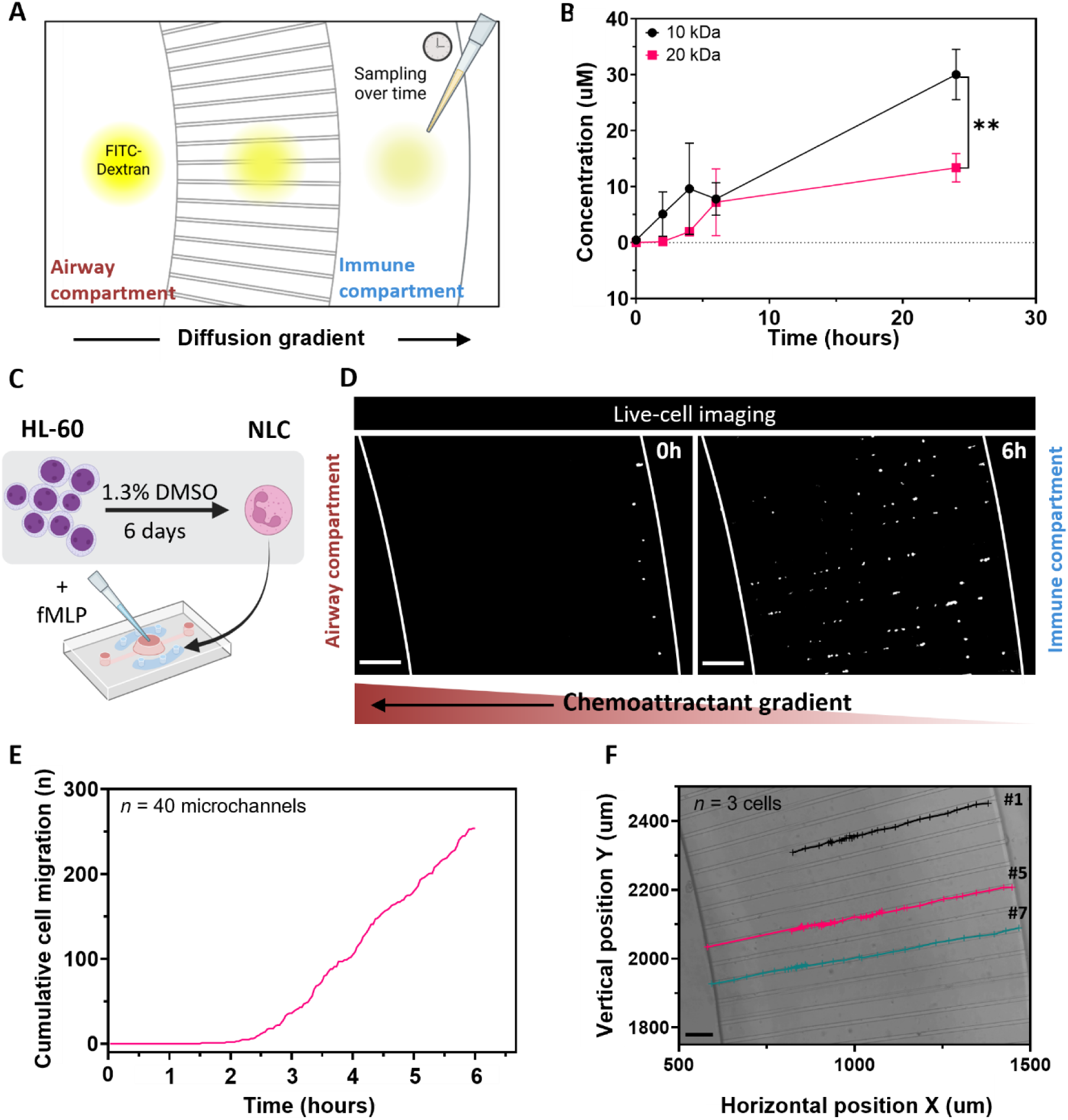
Validation of the microchannel array in scAIR device for soluble factor diffusion and immune cell migration. (A) Schematic of the permeability assay setup used to characterise diffusion across the microchannel barrier in scAIR devices. (B) Quantification of 10 and 20 kDa FITC–dextran concentration in the immune compartment after diffusion across the microchannel array over 24 hours. Data are mean ± s.d, of 3 independent experiments **p < 0.01 (unpaired t-test with Welch’s correction). (C) Schematic of HL-60 differentiation into neutrophil-like cells (NLCs) using 1.3% DMSO, followed by seeding into the immune compartment of the device for fMLP-driven migration. (D) Representative time-lapse images of NLCs labelled with CellTracker Red migration at 0 and 6 hours, showing directional movement of cells across microchannels toward fMLP. Snapshots are from live-cell imaging captured over a 6-hour period. Scale bar represents 100 μm. (E) Manual quantification of the cumulative number of NLCs migrating across the microchannel array to the central airway compartment. NLCs were tracked across n = 40 microchannels in one representative device for a total of 6 hours. (F) Manual tracking of NLC migration across microchannels plotted as XY coordinates to visually represent cell position over time. A representative n = 3 cells tracked across n = 3 microchannels in one scAIR device for a total of 6 hours. Scale bar represents 100 µm.

Next, we assessed whether the generation of a soluble factor gradient across the microchannel array would be at levels conducive to immune cell recruitment. Neutrophil-like cells (NLCs) were differentiated from promyeloblast cells (HL-60) following a well-established protocol ^[31–33]^ (**Figure 3C**), which we validated for the presence of phenotypic neutrophil markers and functional characteristics, including positive CD11b expression and the formation of neutrophil extracellular traps (NETs) (**Supplementary Figure S4**). NLCs were then seeded into the immune compartment of scAIR devices, with simultaneous addition of a potent chemoattractant, N-formylmethionyl-leucyl-phenylalanine (fMLP), into the empty central airway compartment. We leveraged the distinct capability of the scAIR device to capture dynamic NLC migration over a 6-hour live-cell imaging period at single-cell resolution (**Figure 3D, Supplementary Video S1).** Using this dataset, we enumerated NLC recruitment across the microchannel barrier over time, revealing a progressive increase in migrating cells in response to the diffusing chemotactic gradient (**Figure 3E**). By utilising the XY coordinates obtained from manual single-cell tracking, we were able to construct migration trajectories for each migrating cell and overlay these with corresponding microchannels to resolve spatiotemporal migration dynamics at the single-cell level (**Figure. 3F)**. Taken together, these results highlight the capacity of the scAIR device to interrogate immune cell behaviour at single cell level and generate rich trajectory data that can be further leveraged for more comprehensive quantitative analysis.

### 2.3. Machine learning-based analysis automates and resolves heterogeneity in immune cell migration characteristics

While the microchannel array enabled real-time visualisation of immune cell migration at single-cell resolution, extracting meaningful quantitative metrics from these datasets presents a significant analytical challenge. Manual or semi-manual tracking approaches are labour-intensive, prone to user bias, and have limited throughput, thereby restricting analysis to small cell populations and simplified endpoints. Such analyses are insufficient to capture the multidimensional features of single-cell migration trajectories, including speed, persistence, directionality, and path variability, that could define functional heterogeneity among immune cell population.

To address these analytical limitations, we applied a previously developed machine learning analysis (MLA) pipeline ^[34]^, incorporating a state-space tracking model to reconstruct single-cell migration trajectories and enable multi-dimensional feature extraction. Live-cell imaging data were analysed using a structured four-step workflow: (i) microchannel region-of-interest (ROI) definition, (ii) image pre-processing, (iii) cell identification and tracking, and (iv) statistical analysis (**Figure 4A**). Using this framework, microchannel regions were isolated from brightfield images and cells were segmented from fluorescence data following pre-processing to enhance detection of individual NLCs. A state-space model was then used to assign persistent identities across time, enabling reconstruction of continuous single-cell trajectories throughout the imaging period. To ensure robustness of trajectory assignment, model performance was evaluated across key tracking parameters governing detection linking, track initiation, and track termination, which collectively influence whether cell trajectories are merged or fragmented across frames. We optimised the reliability of cell tracking across datasets by systematically varying these key tracking parameters over 100 combinations and then applying a trajectory-based consensus analysis to identify the physical cells reliably detected across models (**Supplementary Figure S5**). To validate accuracy, we benchmarked migration metrics derived from the MLA models against manually annotated ground-truth trajectories generated by two independent annotators. Rather than comparing raw XY positional coordinates, we focused on three core descriptors of migration behaviour: speed, capturing the magnitude of cell movement; accumulated distance, reflecting the total path length travelled; and Euclidean distance, defined as the net distance between initial and final cell positions within the microchannel. The MLA-derived metrics showed strong agreement with manual annotations, enabling robust and unbiased quantification of single-cell migration dynamics (**Figure 4B**).

**Figure 4.**
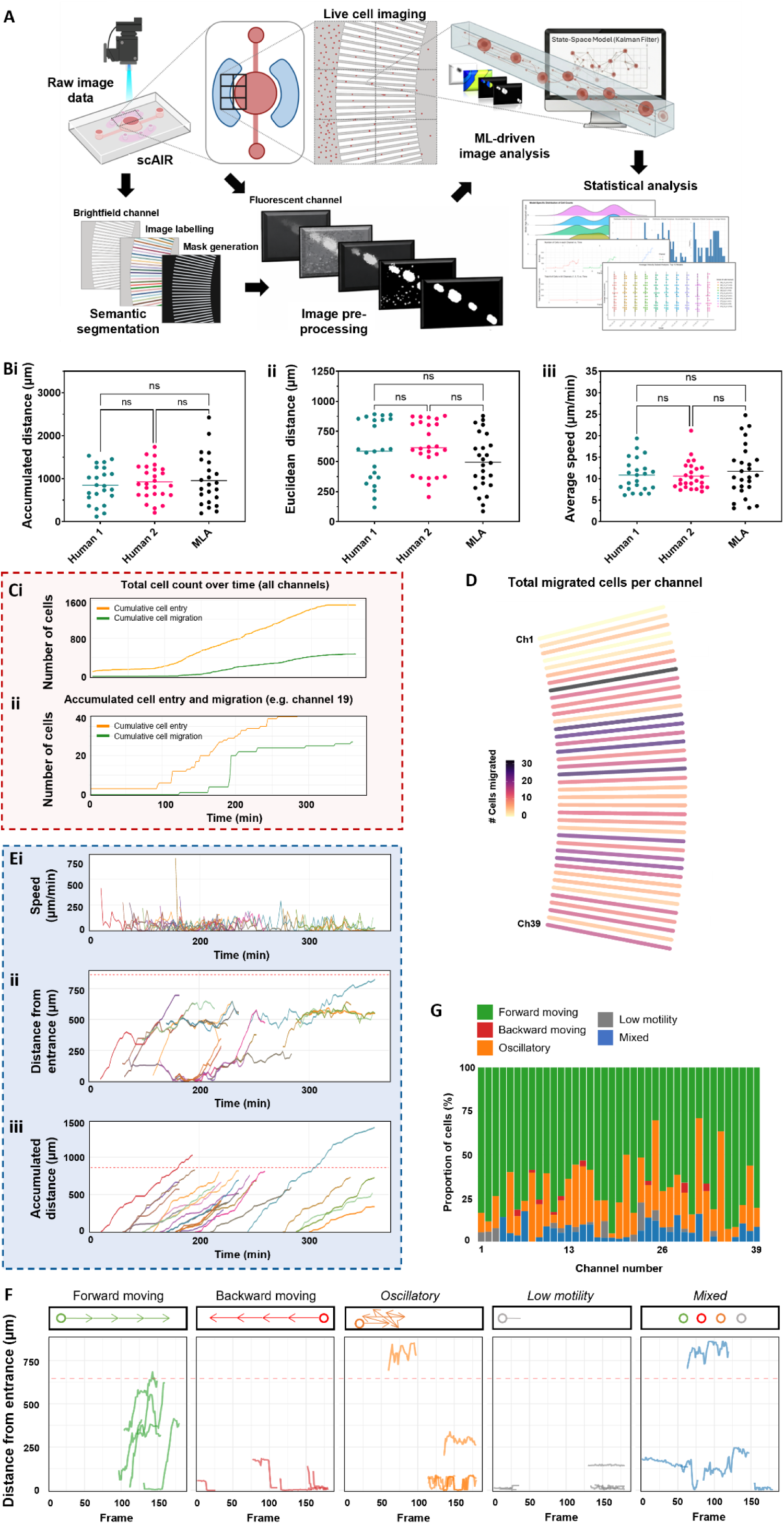
Automated MLA-based assessment of immune cell migration within the microchannel array of scAIR devices. (A) Schematic workflow of the in-house MLA pipeline used to automate immune cell tracking from real-time imaging data. The MLA pipeline is divided into four steps: (i) microchannel region-of-interest (ROI) definition, (ii) image pre-processing, (iii) cell identification and tracking, and (iv) statistical analysis. (B) Comparison of migration metrics obtained using manual tracking by two independent users (green and pink) and the MLA pipeline (black), including (i) accumulated distance, (ii) Euclidean distance, and (iii) speed. Each point represents one cell trajectory and represents the average value between consecutive time frames for single cell trajectories, with the solid line indicating the overall mean. (n = 23-26 cells tracked from one experiment). ns: no statistical significance (One-way ANOVA). (C) MLA enumeration of NLCs across (i) the entire microchannel array, and (ii) within a single representative microchannel. The orange line represents the cumulative number of cells that entered the microchannels over time, while the green line represents the cumulative number of cells that completely migrated across the entire length of the microchannels. (D) Spatial mapping of migration events across the microchannel network using MLA. Each bar represents one microchannel in its relative position in the device. Colour scale from white (0) to black (30) indicates the total number (#) of NLCs fully migrating across the specific microchannel. (E) MLA-generated outputs derived from a single representative microchannel illustrating (i) speed, (ii) distance from entrance (Euclidean displacement), and (iii) accumulated distance. Each coloured line represents a single cell within a representative microchannel. (F) Euclidean distance trajectories of n=4 cells from a single representative microchannel showing different migratory phenotypes, including strong forward, strong backward, oscillatory, and low motility phenotypes. (G) Relative abundance of fMLP-responsive NLCs classified according to distinct migratory phenotypes (n = 1498 cells classified across 39 microchannels).

Finally, we applied the optimised MLA pipeline to resolve single-cell migration behaviours across the microchannels at high spatio-temporal resolution. The algorithm reconstructed individual cell trajectories over the imaging period, enabling quantification of both cumulative and per-cell migration metrics. We enumerated NLCs that had completed migration across the entire microchannel array, as well as those still within individual channels (**Figure 4C (i-ii)**), revealing spatial heterogeneity, non-uniform channel utilisation, and preferential migration paths across the device (**Figure 4D**).

Beyond extracting single-cell endpoint metrics, such as average speed, Euclidean displacement, and accumulated distance shown in Figure 4B, the MLA pipeline also enabled temporal analysis of these behaviours over the course of migration (**Figure 4E, Supplementary Figure S6),** revealing dynamic changes in immune cell motility that would be difficult to capture using conventional manual tracking approaches. Analysis of temporal changes in cell speed revealed that migration occurred in a pulsatile manner rather than at a constant rate, with repeated bursts of movement interspersed with periods of reduced motility **(Figure 4E(i))**. Although NLCs exhibited an average migration speed of 12 ± 6 μm/min **(Figure 4B(iii))**, consistent with values reported for neutrophil migration in comparable microfluidic devices ^[35]^, single-cell tracking demonstrated transient bursts of movement reaching 20–60 μm/min. This observation was consistent with previous studies that have described a similar ‘stop-and-go’ mode whereby immune cells alternate between phases of rapid advancement and reduced motility. ^[36–38]^ Such behaviour has been theorised to improve the efficiency of pathogen localisation within tissue microenvironments. Notably, these transient states are largely obscured when migration tracking is reduced to averaged endpoint measurements alone and were not apparent prior to single-cell trajectory analysis.

Temporal analysis of single-cell Euclidean displacement (net distance from microchannel entrance) and accumulated distance (**Figure 4E (ii–iii), Supplementary Figure S6**) revealed multiple, functionally distinct migration phenotypes within the NLC population. These trajectories encompassed forward migration toward the epithelial compartment, reverse migration, oscillatory movement with repeated directional switching, and low-motility behaviours with minimal net displacement (**Figure 4F**). To quantitatively differentiate this behavioural diversity, we further established three additional quantitative metrics: net progress, tortuosity, and range explored (**Supplementary Table S4**). These parameters enabled automated assignment of individual cells into different migratory phenotypes and quantify their relative abundance within the population (**Figure 4G**). Together, these findings demonstrate that the MLA pipeline can uncover previously unresolved heterogeneity in immune migration dynamics within a single inflammatory microenvironment.

### 2.4. The scAIR recapitulates RSV-driven immune cell recruitment and reveals changes in migratory behaviour with monoclonal antibody treatment

Having demonstrated that the scAIR device supported AEC culture and enabled single cell quantification of immune cell migration in response to exogenous stimuli, we next endeavoured to model real-time crosstalk between immune cells and AECs in response to viral-induced inflammation. By harnessing the MLA-integrated scAIR platform, we sought to define how epithelial-derived inflammatory signalling shapes immune cell recruitment, resolving differences in migratory behaviour under targeted cytokine perturbation.

Upon infection with respiratory syncytial virus (RSV), we found that AECs exhibited a peak level of infection at day 5 post infection (p.i.), with no further increase in RSV N protein mRNA expression at day 7 p.i. (**Supplementary Figure S7**). Since this infection timeline agreed with published ranges of peak infection *in vitro* ^[39]^, we moved forward with a timeline where pre-differentiated AECs were transferred to scAIR devices and immediately infected with RSV for a total of 5 days (**Figure 5A)**. To maximise the concentration of chemokines and cytokines secreted by AECs in the central airway compartment, we established a medium replenishment schedule that preserved AEC viability while promoting the accumulation of soluble factors (**Figure 5A**, **Supplementary Figure S8**). We found that replenishing media daily during the first 3 days p.i., followed by a 48-hour non-replenishment period, allowed for the accumulation of epithelial-derived inflammatory signals within the basal compartment (**Figure 5B, Supplementary Figure S9).** We observed elevated levels of pro-inflammatory cytokines interleukin (IL)-6, IL-8, and tumour necrosis factor-α (TNF-α) (**Figure 5B (i))**, while the secretion of typical type-2 inflammatory cytokines, IL-4, IL-5, and IL-13, was limited or absent. (**Figure 5B(ii))**. These results indicated that RSV infection of AECs from healthy donors in the scAIR produced a predominantly type-1 inflammatory environment, consistent with responses reported following RSV infection in healthy adults. ^[40, 41]^

**Figure 5.**
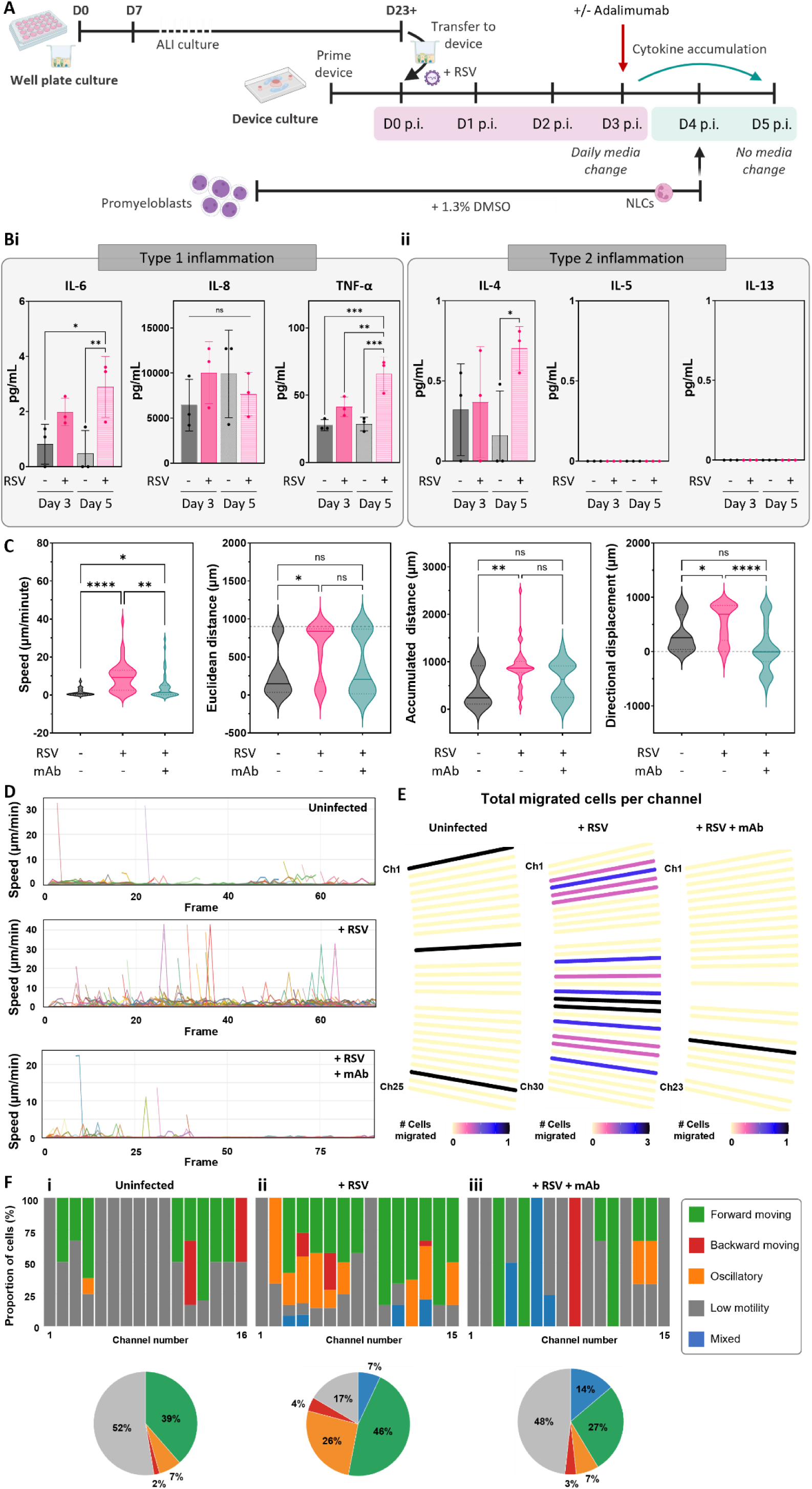
RSV infection of airway epithelial cells induces cytokine secretion and directional immune cell migration. (A) Experimental timeline. AECs were pre-differentiated in Transwell inserts, transferred to scAIR devices, and infected with RSV. Where indicated, the TNF-α-neutralising monoclonal antibody adalimumab was added on day 3 post-infection (p.i.). Media were not replenished between days 3 and 5 p.i. to permit accumulation of basolaterally secreted factors. NLCs were introduced into the immune compartments on day 4 p.i., followed immediately by live-cell imaging. (B) Cytokine and chemokine concentrations in basolateral conditioned media collected on days 3 and 5 p.i. (i) type 1-associated and (ii) type 2-associated factors. Data are presented as mean ± s.d. from n = 3 independent devices. p values were determined by two-way ANOVA with multiple-comparisons correction: ns, not significant; *p < 0.05; **p < 0.01; ***p < 0.001. (C) Violin plots showing single-cell migration parameters, including mean speed, Euclidean displacement, accumulated distance, and directional displacement, for NLCs migrating toward uninfected AECs, RSV-infected AECs, or RSV-infected AECs treated with adalimumab. n = 19 (uninfected), n = 49 (+ RSV), and n = 43 (+ RSV + mAb) cells tracked across 2 independent experiments per condition. Statistical significance was determined using a One-way ANOVA with multiple comparisons. ns: not significant, *p < 0.05, **p < 0.01, ****p < 0.0001. (D) Single-cell migration speeds derived from the MLA pipeline across the complete microchannel array for each experimental condition. Each coloured line represents the trajectory of an individual NLC. (E) Spatial distribution of NLC migration across the microchannel array. Each rectangle represents an individual microchannel at its corresponding spatial position, with colour scale indicating the number of NLCs completing migration through that channel. n = 23-30 microchannels across n = 1-2 independent experiments. (F) Classification of individual NLC trajectories into distinct migratory phenotypes based on MLA-derived trajectory metrics. Bars show the distribution of migratory phenotypes within individual microchannels (top), and corresponding pie charts show their relative abundance across all analysed cells for each experimental condition (bottom). n = 57 (uninfected), n = 115 (+ RSV), and n = 29 (+ RSV + mAb) cells tracked across 1 independent experiment per condition.

We next leveraged the scAIR platform to interrogate neutrophil recruitment in response to an RSV-induced type 1 inflammatory airway microenvironment. NLCs were introduced into the immune compartments at day 4 p.i., coinciding with peak cytokine accumulation, and their migratory dynamics were monitored by live-cell imaging over a 12-hour period (**Supplementary Movies S2–5**). Population-level analysis revealed that RSV infection markedly enhanced NLC recruitment toward the central airway compartment, characterised by increased migration velocity, positive directional displacement, and greater Euclidean and accumulated distances **(Figure 5C)**. Given the marked increase in TNF-α secretion following RSV infection (**Figure 5B (i)**), and its established role as a central regulator of NF-κB-mediated inflammatory signalling, we also investigated whether neutralisation of TNF-α could attenuate the resulting migratory response **(Supplementary Movies S6–7**). Indeed, the addition of adalimumab on day 3 p.i. prior to NLC addition showed significantly reduced migration velocity and directional displacement, while exerting comparatively modest effects on total migration distance, suggesting that adalimumab disrupted chemotactically directed migration toward the infected epithelium without fully suppressing basal motility (**Figure 5C**).

To resolve the cellular mechanisms underpinning these population-level responses, we next examined the spatiotemporal dynamics of individual NLC trajectories within single microchannels. Consistent with bulk recruitment measurements, RSV infection increased the number of migrating cells per channel, whereas uninfected conditions and the addition of adalimumab showed lower number of migrating NLCs (**Supplementary Figure S10)**. Interestingly, we also observed pulsatile migratory behaviour of NLCs towards RSV-infected AECs, with frequent bursts of high-speed movement reaching speeds of 40 μm/min (**Figure 5D (ii))**. In contrast, cells within the uninfected and adalimumab-treated conditions displayed predominantly low-amplitude speed fluctuations, with few transient bursts and overall lower variation in migration speed (**Figure 5D (i, iii))**. Channel occupancy analysis revealed a more pronounced preference for specific microchannels during RSV-induced migration than during fMLP-driven migration (**Figure 4D**), producing a highly heterogeneous spatial pattern across the array (**Figure 5E(i–iii)).** Whereas only three microchannels supported single-cell migration under uninfected conditions and one following adalimumab treatment, RSV infection induced migration across most channels, with multiple NLCs concentrated within selected channels while adjacent channels remained sparsely occupied.

Finally, we classified individual migration trajectories into discrete migratory phenotypes based on net displacement, directional bias, and tortuosity (**Figure 5F (i-iii)**). Under uninfected conditions, migration was dominated by low-motility cells (52%), with the remaining population comprising predominantly forward-moving cells (39%) and only small proportions of oscillatory and backward-moving phenotypes. In contrast, RSV infection shifted the migratory profile towards active recruitment, with forward migration representing the predominant phenotype (46%), accompanied by a marked increase in oscillatory behaviour (26%) and a corresponding reduction in low-motility cells. The emergence of a larger oscillatory population may reflect increased exploratory or less efficiently directed movement following inflammatory activation, whereas the persistence of a forward-migrating subset indicates enhanced directional recruitment capacity. Following TNF-α neutralisation, the phenotypic distribution shifted towards a less migratory state, with low-motility cells again representing the largest population (48%), while the proportion of forward-moving cells decreased to 27%. The persistence of backward and mixed trajectories (14%) despite restoration of a predominantly low-motility population indicates that TNF-α neutralisation does not simply return neutrophil behaviour to baseline, but instead produces a distinct migratory state characterised by reduced forward movement and increased trajectory variability. Collectively, these results demonstrate that the MLA-integrated scAIR platform can resolve subtle yet functionally distinct immune migration behaviours that are obscured by conventional population-averaged assays.

## 3. Discussion

The dynamic interplay between airway epithelial and immune cells is a key determinant of variability in disease progression and therapeutic response during airway infection, yet it remains difficult to investigate using existing in vitro models. Current airway-immune co-culture systems typically integrate epithelial cell maturation and immune cell compartmentalisation within a single engineered construct, such as hydrogel barriers or membrane-based devices, making it challenging to temporally coordinate the sequential processes of pathogen sensing, epithelial activation, and immune cell recruitment that occur during airway infection. To address this limitation, we developed the scAIR platform, which deliberately decouples epithelial maturation from immune interaction assays by culturing differentiated airway epithelium on removable Transwell inserts while independently controlling immune compartmentalisation through a microchannel interface. Paired with single-cell MLA, this platform enabled quantitative interrogation of immune behaviours that are obscured by conventional population-averaged measurements. Using RSV-induced inflammation as a proof-of-concept model, we demonstrated that the scAIR platform not only recapitulates NLC recruitment toward an inflamed airway epithelium but also resolves previously unrecognised behavioural heterogeneity and differential responses to inhibition of TNF-α signalling. Together, these findings highlight the value of combining modular airway–immune co-culture with high-content single-cell imaging analytics to uncover mechanisms governing immune recruitment and function in complex inflammatory microenvironments.

Successful modelling of airway infection requires a mature epithelium that faithfully recapitulates key physiological functions, including barrier integrity and mucociliary activity. Many respiratory viruses exhibit cell-type tropism and preferentially infect specialised epithelial populations, such as ciliated cells, that emerge only after full differentiation and express the receptors required for viral attachment and entry. ^[42]^ Furthermore, an intact epithelial barrier is essential for evaluating infection-induced barrier disruption, inflammatory responses, and therapeutic efficacy. While the scAIR device was specifically designed to interrogate epithelial-immune crosstalk, we acknowledge that it represents a simplified system compared with advanced lung-on-chip (LOC) platforms that incorporate perfusion culture and/or cyclic mechanical actuation to mimic airway biomechanics ^[12]^. Although these dynamic features have been reported to further enhance epithelial differentiation and mucociliary maturation ^[43–45]^, well-established static ALI cultures already achieve a high degree of physiological functionality, including robust barrier integrity, ciliation, mucus production, and viral susceptibility ^[15, 46–48]^, which was also demonstrated in our study (**Figure 2**). As such, dynamic biomechanical stimulation may not be critical for modelling epithelial-driven immune recruitment in the context examined here. Importantly, the platform supported RSV-induced inflammatory signalling sufficient to drive NLC recruitment **(Figure 5B (i-ii))** and enabled quantification of the attenuated immune responses following monoclonal antibody treatment **(Figure 5C-F)**. Thus, by decoupling epithelial maturation from immune interaction assays, the scAIR platform preserves the robustness and accessibility of static ALI culture while enabling controlled and quantitative investigation of airway–immune crosstalk.

Immune cell recruitment to inflammatory sites is often inferred indirectly by quantifying cytokine production or the number of immune cells reaching the site of injury. However, growing evidence from intravital imaging has revealed that immune recruitment is fundamentally a dynamic behavioural process, comprising heterogeneous migratory phenotypes that continuously adapt to local inflammatory cues. ^[49–51]^ Capturing this behavioural diversity is therefore critical for understanding how immune responses contribute to disease progression and therapeutic efficacy. Although many in vitro airway models faithfully recapitulate epithelial differentiation and immune–epithelial interactions, their analytical outputs remain largely restricted to population-level measurements, such as cytokine production or the total number of recruited cells. This is largely because robust quantification of single-cell migration across large live-cell imaging datasets remains technically challenging. Manual and semi-automated tracking approaches can extract detailed migration trajectories ^[23, 52–54]^, but they are labour-intensive, susceptible to user bias, and impractical for analysing the thousands of cells generated during long-term live-cell imaging.

MLA has recently emerged as a powerful approach for quantifying cell motility and chemotactic responses in simplified microfluidic migration assays. ^[55]^ However, its application to physiologically relevant tissue models have remained limited by device architecture. For example, vertically stacked coculture systems place the responding and target cell populations on different imaging planes, making continuous uninterrupted tracking of the entire recruitment process challenging. ^[13, 18, 56]^ Conversely, devices designed for high-content migration analysis frequently lack the tissue context required to investigate immune–epithelial interactions. ^[34, 57–59]^ To overcome these limitations, we deliberately designed the scAIR platform and MLA pipeline as an integrated experimental–computational framework. The horizontally compartmentalised architecture enables continuous imaging of immune cell recruitment toward an inflamed airway epithelium within a single focal plane, while the MLA automatically reconstructs thousands of individual trajectories over extended imaging periods and extracts multidimensional behavioural descriptors from each migrating cell. Rather than simply increasing analytical throughput, this integrated framework expands the biological information that can be obtained from immune recruitment assays. Beyond quantifying the number of recruited cells, the scAIR platform resolves immune recruitment as a dynamic behavioural process, revealing how individual migratory phenotypes emerge, evolve, and respond to therapeutic intervention.

Using a state-space tracking model, MLA-driven image analysis automatically reconstructed trajectories for large numbers of immune cells migrating across multiple microchannels over an 8-hour period in response to epithelial-derived chemotactic signals. Importantly, the robustness of automated tracking depended on careful optimisation of trajectory assignment parameters to balance track fragmentation against erroneous track merging. In comparison, earlier cell tracking approaches typically relied on nearest-neighbour or centroid-linkage algorithms, which can become unreliable in the presence of missed detections, cell proximity, or complex migratory trajectories. ^[60, 61]^ More recently, deep learning-based tracking frameworks combining convolutional neural network-driven cell detection with trajectory reconstruction have demonstrated improved performance in dense and heterogeneous imaging datasets. ^[62]^ However, the Kalman filter–based state-space model employed here was particularly well suited to our application because cells migrated within geometrically constrained microchannels, where future positions could be predicted with reasonable accuracy. ^[63]^ This enabled robust trajectory reconstruction while maintaining computational efficiency and avoiding the substantial training data requirements associated with deep learning approaches.

A central question raised following RSV infection in the scAIR platform is whether immune cell recruitment occurs independently between cells, or whether the migration of one cell can influence the behaviour of the larger population. Emerging work on migrasomes and neutrophil-derived migratory trails suggests that migrating immune cells can deposit extracellular signals that influence surrounding cell behaviour. ^[64, 65]^ This provides a potential explanation for the preferential use of individual migration tracks used by NLCs responding to infection-driven cues **(Figure 5E)**. Although the present study cannot determine whether these mechanisms contribute to the preferential track usage observed during RSV infection, the spatial pattern is consistent with the possibility that migrating NLCs influence subsequent recruitment through localised signals. The distribution of migratory phenotypes also suggests that RSV infection alters the composition of the migrating NLC population, rather than simply increasing the overall extent of migration. The predominance of forward-migrating phenotypes following RSV infection, compared with the low-motility population observed in the uninfected condition, is consistent with inflammatory cues shifting neutrophils between distinct migratory states. ^[66]^ The spatial distribution of these behaviours is therefore important when interpreting the effect of TNF-α neutralisation. Although the population-level change in directional migration could suggest that TNFα blockade broadly redirects NLC chemotaxis, individual channel-resolved analysis indicates that the apparent shift was concentrated within a single migration path rather than representing widespread reversal across the device **(Figure 5F(iii))**. This may reflect the integration of locally varying chemotactic signals, as neutrophils respond to combinations of competing chemoattractants rather than to a single uniform directional cue. ^[67, 68]^ The biological interpretation therefore suggests that TNF-α neutralisation did not simply reverse NLC migration but altered recruitment within a spatially restricted region of the device. This may reflect the broader role of TNFα in regulating neutrophil recruitment indirectly through the inflammatory microenvironment, including involvement of NF-κB signalling, rather than acting as a direct chemoattractant. ^[69, 70]^ These findings show that inflammatory conditions can alter both the behavioural composition and spatial organisation of NLC recruitment, rather than simply changing the overall number or direction of migrating cells.

In summary, scAIR establishes a versatile human-relevant platform for interrogating epithelial–immune crosstalk through dynamic, multiparametric analysis of immune cell behaviour at single-cell resolution. By combining physiologically relevant airway biology with automated trajectory analysis, the platform transforms immune recruitment from a simple endpoint measurement into a quantitative functional readout of inflammatory responses. Although demonstrated here using RSV infection and TNF-α blockade, the modular design readily accommodates diverse epithelial and immune cell populations, enabling mechanistic studies of disease-specific inflammation, systematic evaluation of immunomodulatory therapies, and investigation of patient-to-patient variability using primary or patient-derived cells. As human-relevant microphysiological systems continue to emerge as New Approach Methodologies (NAMs) for preclinical research, the scAIR platform provides a scalable framework for uncovering inflammatory endotypes, accelerating therapeutic discovery, and advancing precision models of respiratory disease.

## 4. Experimental Section

### 4.1. Fabrication of the scAIR device

#### 4.1.1. scAIR device design and fabrication

A negative photoresist of the microfluidic layer was designed using AutoCAD 2022 (Autodesk Inc., United States), and photolithography was used to generate a master mould. Briefly, negative photoresists (Mechanobiology Institute, Singapore) were patterned on a 4” silicon wafer to create a mould for airway compartments 100 µm in height, and microchannels 5 µm in height. To increase throughput and reproducibility, negative moulds of the reservoir layers were designed using SolidWorks 2022 (Dassault Systèmes, France) and printed with Moiin HighTemp resin (DMG Enterprises SE, Germany) on the ASIGA Max X27 DLP 3D printer (ASIGA, Australia) with a 0.1 mm layer height. Post-printing, the moulds were rinsed in 100% isopropanol (IPA; Sigma Aldrich, Australia) for 10 minutes, followed by a UV cure for 2 hours at 40°C. The SU-8 master mould and negative reservoir layer moulds were exposed to trichloro(1h,1h,2h, 2 h) perfluorooctyl silane (Sigma-Aldrich, Australia) treatment and rinsed with 100% IPA prior to use. Polydimethylsiloxane (PDMS) was prepared using a 10:1 (w/w) ratio suspension of SYLGARD™ 184 polymeric base and curing agent (Dow Chemical Company, Australia) and poured onto the negative reservoir moulds and degassed for 1 hour. Plastic spacers 100 µm in height were placed on either side of the SU-8 master mould and PDMS was carefully poured to overfill the spacers and degassed for 1 hour. To create a uniform height, an OHP transparency sheet (Suremark, Singapore) was placed on top of the SU-8 and reservoir layer moulds, topped with a metal block to apply uniform weight and cured at 70°C for 4 hours.

#### 4.1.2. scAIR device assembly

Cured PDMS layers were peeled from their respective moulds using flat tweezers and cut to shape with a scalpel. Through holes and inlets were punched into the PDMS replicas using a biopsy punch (Miltex, United States) before being rinsed with 100% IPA and distilled water and dried using a handheld air pump. Devices were assembled by exposing PDMS or glass components to a 5:15 ratio of oxygen and argon, respectively, for 60 seconds at 30 W RF power (Expanded Plasma Cleaner, Harrick Plasma, United States). Briefly, the top PDMS reservoir layer was irreversibly bonded to the middle reservoir layer, forming the reservoir complex, which was then bonded to the bottom microfluidic layer. Finally, the entire PDMS fluidic-reservoir complex was bonded to 75 mm x 25 mm (l x w) glass slides and placed between two plates at 70°C for 2 h to ensure sufficient bonding. Prior to any cell experiments, devices were sterilised with 70% v/v EtOH for 1 hour, dried at 70°C, and rinsed with 1X PBS for a minimum of 2 hours.

### 4.2. Diffusion study of fluorescent tracers in scAIR devices

Devices were sterilised with 70% v/v EtOH and dried at 70°C before being rinsed and primed with phosphate-buffered saline (PBS). Solutions of 100 μM 10 and 20 kDa FITC-dextran (Thermo Fisher Scientific, Australia) were added to the airway compartment of devices and allowed to diffuse across the microchannels into the immune compartments. Samples of 25 μL were collected from both immune compartments immediately at 0 hours and read in a microplate reader (CLARIOstar Plus, BMG Labtech, Australia) at ex 490 em 520, then returned to the same immune compartment to maintain volume. Sample collections and readings were repeated every 2 hours up to hour 6, then a final sample was taken at 24 hours. Diffused 10 and 20 kDa FITC-dextran concentrations were calculated by interpolating fluorescence intensities against a standard curve prepared from known FITC-dextran concentrations at each timepoint.

### 4.3. Cell culture

#### 4.3.1. Airway epithelial cell culture

Primary human bronchiolar epithelial cells (HBEC; Lonza, Switzerland) were cultured in PneumaCult^TM^-Ex Plus media (PEPM) supplemented with PneumaCult^TM^-Ex Plus 50X Supplement, 96 ng/mL hydrocortisone and 1% penicillin/streptomycin (Sigma Aldrich, United States). Media was refreshed every 48 hours until cells reached 70% confluency, then HBECs were plated at a density of 5×10^4^ cells/well onto the apical side of Transwell^®^ inserts with 0.4 µm polyester membranes coated with 0.05 mg/mL collagen I rat tail (Enzo Life Sciences, United States). The basal chamber was filled with 600 μL of PEPM and media was replenished every 48 hours until cells reached 100% confluency. At this point, apical media was completely removed to initiate ALI culture, and basal media was replaced with PneumaCult^TM^-ALI Maintenance Media (PAMM) supplemented with 10% PneumaCult^TM^-ALI 10X Supplement, 1% PneumaCult^TM^-ALI Maintenance Supplement, 4 µg/mL heparin solution, 480 ng/mL hydrocortisone stock solution and 1% penicillin/streptomycin (Sigma Aldrich, United States). Basal media was replenished every 48 hours until cells were fully differentiated and ready for further assays. Apical surfaces were washed once per week with 1X PBS to remove excess mucus.

#### 4.3.2. Immune cell culture

Promyeloblast cells (HL-60; ATCC, United States) were expanded in suspension culture in Iscove’s Modified Dulbecco’s Medium (IMDM; Thermo Fisher Scientific, Australia) supplemented with 20% FBS (Thermo Fisher Scientific, Australia) and 1% penicillin/streptomycin (Sigma Aldrich, United States) and maintained at a cell density between 1×10^5^ and 1×10^6^ cells/mL. HL-60 differentiation into neutrophil-like cells (NLCs) was achieved using previously established protocols ^[31–33]^. Briefly, HL-60 cells were cultured in IMDM containing 1.3% dimethyl sulfoxide (DMSO; Thermo Fisher Scientific, Australia), 0.5% FBS (Thermo Fisher Scientific, Australia) and 1% penicillin/ streptomycin (Sigma Aldrich, United States) for 6 days without media replenishment. On day six, NLCs were resuspended in complete IMDM growth medium and prepared for further assays.

### 4.4. Cell viability assay

AEC cultures were treated with 2µg/mL fluorescein diacetate (FDA; Thermo Fisher Scientific, United States) and propidium iodide (PI; Thermo Fisher Scientific, United States) apically for 10 minutes at 37 °C, then counterstained with 1µg/mL Hoechst 33342 (Thermo Fisher Scientific, United States) for 5 minutes at room temperature. Stained cells were imaged on a fluorescent microscope (Zeiss Axio Imager 2.0). Cell viability was quantified and results expressed as percentage cell viability, calculated by dividing the number of live cells by total cells and multiplying by 100, and plotted using GraphPad Prism software.

### 4.5. TEER measurement

Barrier integrity was assessed by measuring transepithelial electrical resistance (TEER) of the epithelium using an epithelial voltohmmeter (World Precision Instruments, United States). AEC cultures were assessed for barrier integrity prior to device insertion and after 5 days of device culture. Transwell^®^ inserts that had been in device culture were transferred back to 24 well plates to perform the TEER assay. TEER was measured by placing the electrode probes in PBS-containing apical and basolateral chambers, and the resistance recorded in ohms (Ω). The resistance of a Transwell^®^ membrane void of cells was recorded as a blank control at the time of each measurement. TEER was calculated according to the following equation:

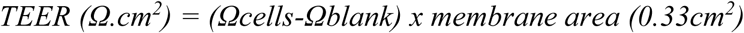

### 4.6. Immunofluorescence

At room temperature, AEC cultures were washed twice with PBS, fixed with 4% paraformaldehyde (PFA; Sigma Aldrich, United States) for 15 minutes, and washed a further two times in PBS. The Transwell^®^ membrane was carefully cut from the insert using a scalpel and placed in a 24 well plate. Cell-laden membranes were permeabilised using 0.3% Triton X-100 (Sigma Aldrich, United States) for 10 minutes, then blocked with 5% BSA in PBS to prevent non-specific binding. After 2 hours, cells were incubated with primary antibodies MUC5AC (Abcam, United Kingdom), acetylated alpha-tubulin (Abcam, United Kingdom) and ZO-1 (Invitrogen, United States) at 1:250, 1:500 and 1:50 dilutions, respectively, in 5% BSA overnight at 4°C on a plate rocker. The following day, cells were washed three times in PBS and corresponding secondary antibodies were added at a dilution of 1:250 and incubated at room temperature for 2 hours in the dark. Membranes were washed a further three times in PBS before being incubated with 4,6-diamidino-2-phenylindole (DAPI; Thermo Fisher Scientific, United States) at 1:1000 dilution for 10 minutes at room temperature in the dark. Finally, membranes were washed twice in PBS before being mounted to a glass slide and imaged using a confocal microscope (Nikon A1R, Nikon, Japan) at 40X and 63X objectives.

### 4.7. On-device RSV infection of airway epithelial cells

Sterile devices were primed with cell culture medium and incubated at 37°C overnight. Pre-differentiated AECs cultured on Transwell^®^ inserts were depleted of steroids and antibiotics 24 hours prior to infection, then transferred to the airway compartment of each device. Immediately following, 80 µL of RSV inoculum (RSV4; propagated and titrated as previously described ^[71]^) at 1.7×10^6^ pfu/mL was placed on the apical surface of AECs and incubated for 4 hours at 37°C. Uninfected controls were subjected apically to 80 µL of steroid and antibiotic-free PAMM for 4 hours. After this time, ALI culture was re-introduced by removing the apical media, and basal media was replenished with steroid and antibiotic-free complete PAPM via the airway chamber inlets every 24 hours for the first three days. No media change was performed between day 3 and 5 post-infection (p.i.) to allow for cytokine gradient generation. On day 4 p.i., NLCs were labelled with 2.5 µM CellTracker Red (Thermo Fisher Scientific, Australia) and incubated for 30 minutes at 37°C before being resuspended in serum-free IMDM. Labelled cells were introduced to the immune compartments of devices at a cell density of 3×10^6^ cells/mL in a total volume of 20 µL per immune compartment.

### 4.8. Adalimumab treatment

RSV infection of AECs on device was performed as previously described, however on day 3 post-RSV infection, the final media change before NLC introduction consisted of media containing 2.5 ng/mL Adalimumab (Thermo Fisher Scientific, Australia).

### 4.9. Cytokine quantification

Cytokines present in conditioned media from the basal side of infected or uninfected AEC device cultures at day 3 and day 5 post-RSV infection were analysed using the Bio-Plex pro human cytokine 17-plex panel (Bio-Rad, United States) according to manufacturer guidelines. Samples were run in a 96-well plate using the Luminex^®^ system (Bio-Plex^®^-100, Bio-Rad). Samples were run in two technical replicates and three biological replicates from each timepoint.

### 4.10. NLC migration assay in response to fMLP

NLCs were adjusted to a cell density of 4×10^6^ cells/mL in complete IMDM media and seeded into the immune compartments of scAIR devices. After 30 minutes of incubation at 37°C, 5% CO_2_, the central airway compartment was flushed with 500 µL of 50 ng/mL fMLP in serum-free IMDM.

### 4.11. Live-cell imaging of NLC migration

Live-cell imaging was performed using a fluorescent microscope (Leica AF6000 LX) housed in an environmental chamber calibrated to 37°C, 5% CO₂. Images of the device were captured at 10X magnification in brightfield and TRITC (577/602 nm) channels with a tile scan acquisition at specified time points **(Supplementary Table S2)**. Following imaging, datasets were auto stitched and NLC migration parameters were quantified manually using ImageJ or using an in-house MLA pipeline.

### 4.12. Development of machine learning analysis (MLA) pipeline for migration quantification

A MLA computer-vision workflow was developed to assist in the quantification and analysis of recorded imaging data (**Supplementary Figure S1**). This workflow automates the task of having to identify, track and quantify cell positions and movement throughout the live cell imaging experiments and serves to standardise image processing and analytical output with a reproducible and well-documented procedure. The in-house developed workflow involves four key steps: mask generation, image preprocessing, and image analysis, which were performed in MATLAB, followed by statistical analysis conducted in R (R Core Team (2025) Vienna, Austria. Available at: https://www.R-project.org/). Raw image datasets were comprised of stitched 8-bit tag image file (TIF) formatted images for both the brightfield and fluorescent (TRITC; 577/602 nm) channels in a time series acquired using the fluorescent microscope (Leica AF6000 LX) described earlier. Information on the individual datasets is available in **Supplementary Table S2**. The following sections briefly describe key steps of the MLA pipeline:

#### 4.12.1. Mask Generation

Prior to image processing and analysis, a masking step was performed to define microchannel regions of interest (ROIs) within each time-lapse image. Depending on the degree of field-of-view drift observed during the early timepoints of live-cell imaging, one to eight masks were manually generated per dataset using the MATLAB image labelling application and applied to the first several brightfield images. These masks were subsequently propagated across the full time series to ensure consistent identification and localisation of individual microchannels in all frames. Binary representations of the masks were additionally used to define the entry points from the immune compartment and exit points into the airway compartment for each microchannel. Identification of these points was required to account for dataset-specific variations in microchannel orientation, position, and direction of cell migration.

#### 4.12.2. Image pre-processing

Each fluorescent time-lapse image was automatically pre-processed using a standardised pipeline, including contrast adjustment, masking, morphological filtering, thresholding, binarisation, and erosion. Preprocessing parameters were optimised using representative keyframes from each dataset and subsequently applied consistently across the full time series or linearly interpolated over time where appropriate. Specific parameter values for each dataset can be found in **Supplementary Table S3**.

#### 4.12.3. Image analysis

Image analysis was performed on pre-processed, binarized images to identify and quantify individual cells in each frame. For each cell, the dataset recorded a unique ID, the microchannel of residence, centroid coordinates, cell area, distance from the microchannel entrance, accumulated distance travelled, and instantaneous velocity. Segmentation was performed using a Euclidean distance transform followed by watershed separation to distinguish closely spaced or overlapping cells (**Supplementary Figure S2-3**). Cell tracking was implemented using a discrete-time, linear state-space model (Kalman filter) to predict cell positions in subsequent frames. A cost matrix, based on the squared Mahalanobis distance between predicted and observed positions with a log-determinant penalty, was constructed and solved using the Munkres variant of the Hungarian assignment algorithm to associate detections with existing tracks. Non-assignment costs were optimised per dataset to balance track continuity and the creation of new tracks. The outputs of this assignment process, including cell coordinates, assigned track IDs, and unassigned detections, were used to update the Kalman filter and recorded in the main dataset table for each experiment.

#### 4.12.4. Statistical analysis on MLA conditions

All downstream analysis was performed in R. Master data tables containing the frame-by-frame state-space, cell-accumulation, and cell-migration data generated in MATLAB were imported into R and combined into a single long-format data frame. Pixel-based quantities were converted to physical units using the dataset-specific pixel size, and per-frame Euclidean distance from the entrance, accumulated travelled distance, and instantaneous velocity were derived for every track in every model.

### 4.13. Statistical analysis

Numerical data are presented as mean ± standard deviation. All statistical analysis was performed using GraphPad Prism (version 9), excluding any MLA which was performed in R, as detailed above. Replicate number, statistical test type and thresholds for p values are indicted in the corresponding Figure legends.

## Acknowledgements

This work was supported by the Australian Research Council (DP200101658, DP230100721 and IC250100027) and the Max Planck Queensland Centre (324912-007), awarded to Y.-C. T. This work was enabled by the Central Analytical Research Facility (CARF) at the Queensland University of Technology (QUT). We gratefully acknowledge Christina Theodoropoulos for her technical assistance and support. Open access publishing was facilitated by Queensland University of Technology under the Wiley–QUT agreement via the Council of Australian University Librarians.

## Author Contributions

Conceptualisation and experiment designs were performed by L.-M.Y, Y.-C.T. and K.S. Experiments were conducted by L.-M.Y. and J.-Y.K. All machine learning algorithm generation and analysis were performed by C.P.T, L.-M.Y., and J.-Y.K., with support from R.D. Data analysis was performed by L.-M.Y. and J.-Y.K. J.A.C contributed to the design and CAD modelling of the microfluidic device. Writing and editing of the manuscript was performed by L.-M.Y., C.P.T., J.-Y.K., K.S., and Y.-C.T.

## Data Availability Statement

The data that support the findings of this study are available from the corresponding author upon reasonable request.

Received: ((will be filled in by the editorial staff))

Revised: ((will be filled in by the editorial staff))

Published online: ((will be filled in by the editorial staff))

## Table of Contents

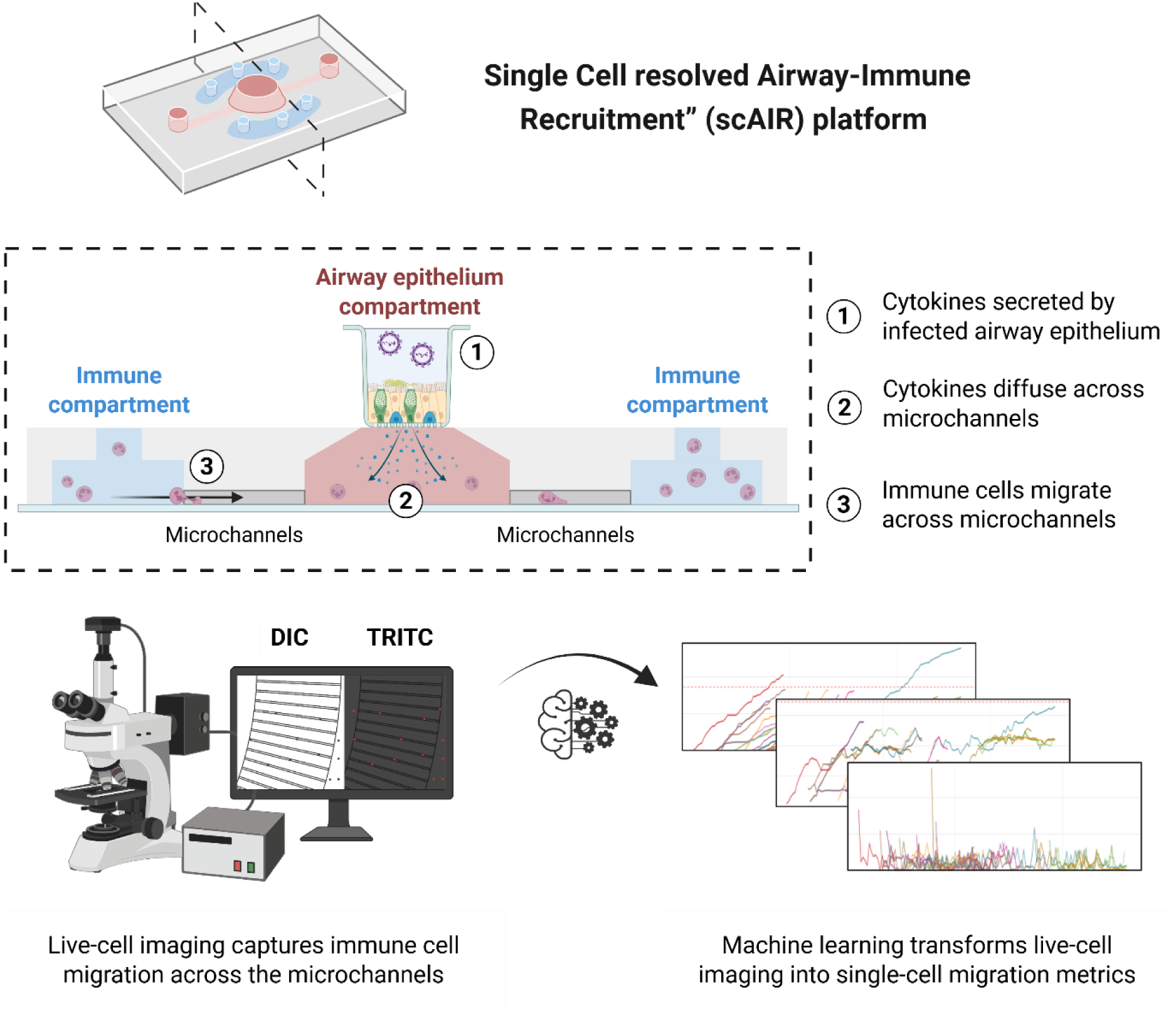

A microfluidic airway model coupled with machine learning that recreates infection-driven immune responses in the lung. The “Single Cell resolved Airway-Immune Recruitment” (scAIR) platform tracks individual immune cells as they move toward infected airway tissue and reveals diverse movement patterns that are obscured in standard assays. It also shows how blocking certain inflammatory factors changes immune cell behaviour during viral inflammation.

## Supporting Information

### Supplementary Method S1: Flow cytometry to confirm NLC differentiation

NLCs were collected from plates and washed three times with 1X PBS to remove residual cell media. Cells were pelleted by centrifugation and resuspended at a density of 1×10^6^ cells/mL in LIVE/DEAD Fixable Viability Dye eFluor™ 520 (Thermo Fisher Scientific, Australia) at a 1:1000 dilution in flow cytometry staining buffer. The cells were allowed to incubate for 30 minutes at 4 °C, protected from light. To block non-specific Fc-mediated interactions, cells were incubated with 1 mg/mL of anti-human Fc receptor binding inhibitor antibody (Thermo Fisher Scientific, Australia) for 15 minutes at room temperature, then incubated with 10 µg/mL of CD11b Monoclonal Antibody PE-Cyanine7 for 30 minutes at 4 °C, protected from light. Flow cytometry was performed immediately using the BD FACSCelesta™ Cell Analyzer (BD Biosciences, United States) and data visualisation/gating was performed via the in-built FACSDiva™ software. After checking fluorochrome voltages with stained and unstained controls, the following settings were used throughout flow cytometry experiments (**Supplementary Table S1**). The PE-Cy7 filter and the BD Horizon Brilliant™ Violet 510 (BV510) filter were used to assess CD11b and viability fluorescence, respectively. FlowJo Software (BD Biosciences, United States) was used to gate and analyse samples, and results were plotted using GraphPad Prism.

### Supplementary Method S2: Manual quantification of immune cell migration

Stitched datasets from live-cell imaging experiments were imported to ImageJ with merged brightfield and TRITC (577/602 nm) channels and cropped to the first 6 hours. Using the inbuilt TrackMate Manual Tracking plugin (ImageJ), NLCs were tracked crossing the microchannels by sequentially clicking on the position of each cell over time. This generated a unique XY coordinate for each cell. Tracking was only completed for cells in the dataset that interacted with the microchannel, regardless of whether they completed the migration to the central airway compartment. Since some NLCs were already within the microchannels at *t=0* of the live-cell imaging experiment, these cells were not used for Euclidean distance and accumulated distance calculations, since their inherent shortened paths may skew the results. The resulting XY data for each cell was converted from pixels to microns, and analysed for NLC migration parameters determined as follows:

Euclidean distance, referring to the direct measurement between the initial position and final position of a cell’s movement only, was determined by **Equation 1**. Where (*^X^start* and *^Y^start*) are coordinates of the cells starting position within the microchannel, and (*^X^end* and *^Y^end*) represent the final coordinates of the cells position.

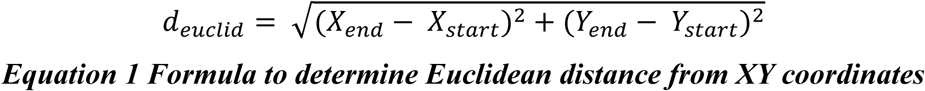

To measure the total path travelled by a cell, accounting for changes in direction, accumulated distance was calculated by summing the distances between consecutive points along the cell’s trajectory using **Equation 2**. Where *d_i_* is the distance in µm between two consecutive points, and *n* represents the total number of points in the path.

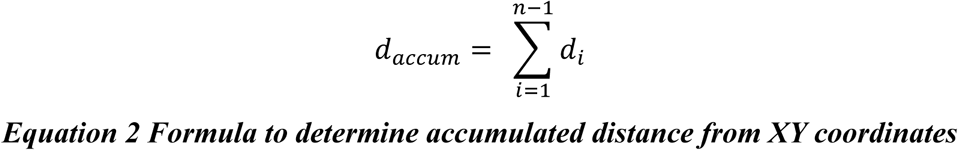

Average speed was calculated as the mean of frame-to-frame speeds obtained from live-cell imaging. For each imaging interval, speed was defined as the distance travelled between consecutive cell positions divided by the time between frames, such that Δ*d_i_* represents the distance travelled between frame *i* and *i* + 1 and Δ*t* represents the imaging interval in minutes (**Equation 3**). Frame-to-frame speeds were then averaged across all intervals for each cell and are reported in µm/min.

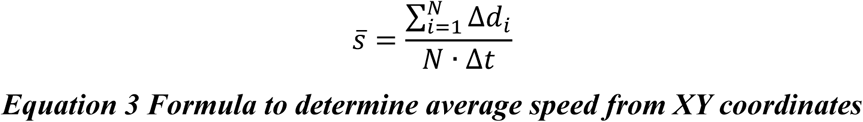

### Supplementary Method S3: RT-qPCR analysis

Total RNA was isolated from HBECs using the RNeasy Mini Kit (Qiagen, Germany) according to manufacturers instructions and quantified using a NanoDropOneC (Thermo Fisher, United States). RNA samples were diluted to a concentration of 10 ng/μL and cDNA synthesis was performed using the SensiFAST cDNA Synthesis Kit (Meridian Bioscience, United States). RT-qPCR was performed using SensiFAST™ SYBR Lo-ROX One-Step Kit (Meridian Bioscience, United States) on a QuantStudio 6 Flex (Thermo Fisher Scientific, United States). Relative gene expression (fold change) was calculated relative to β-actin using the 2-^ΔΔCT^ method. Primer sequences used are outlined in **Supplementary Table S5** (Kicqstart Primers, Sigma Aldrich, United States).

**Supplementary Figure S1.**
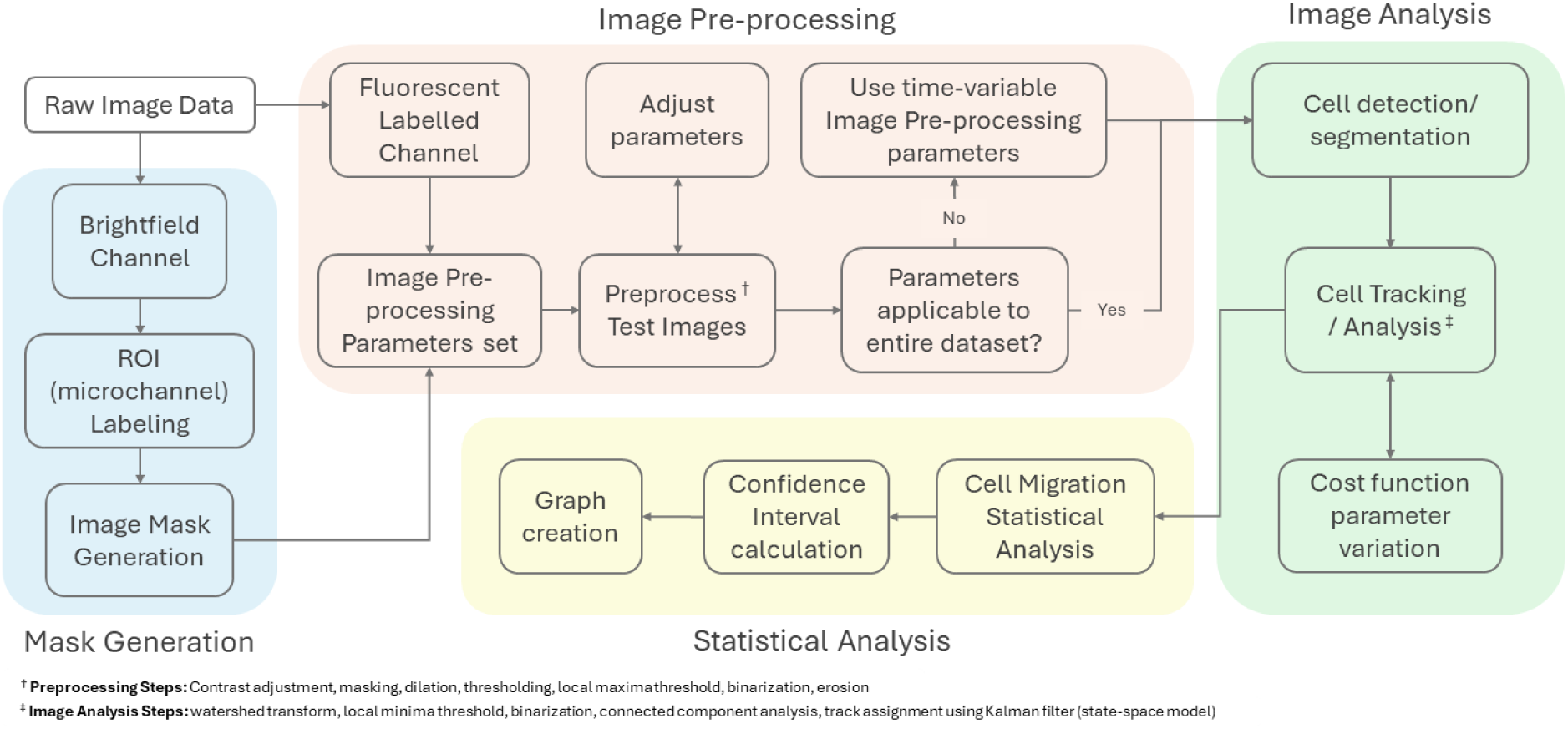
Machine learning (ML)-based workflow for live-cell imaging analysis. Flow chart illustrating the in-house developed workflow used to quantify and analyse recorded live-cell imaging data. The workflow standardises image processing and analytical output through a reproducible and well-documented pipeline that automates cell identification, tracking, and quantification of cell position and movement. The workflow comprises four major stages with associated sub-steps: (1) mask generation, (2) image preprocessing, and (3) image analysis, implemented in MATLAB, followed by (4) statistical analysis performed in R. Each stage and its corresponding sub-processes are shown schematically.

**Supplementary Figure S2.**
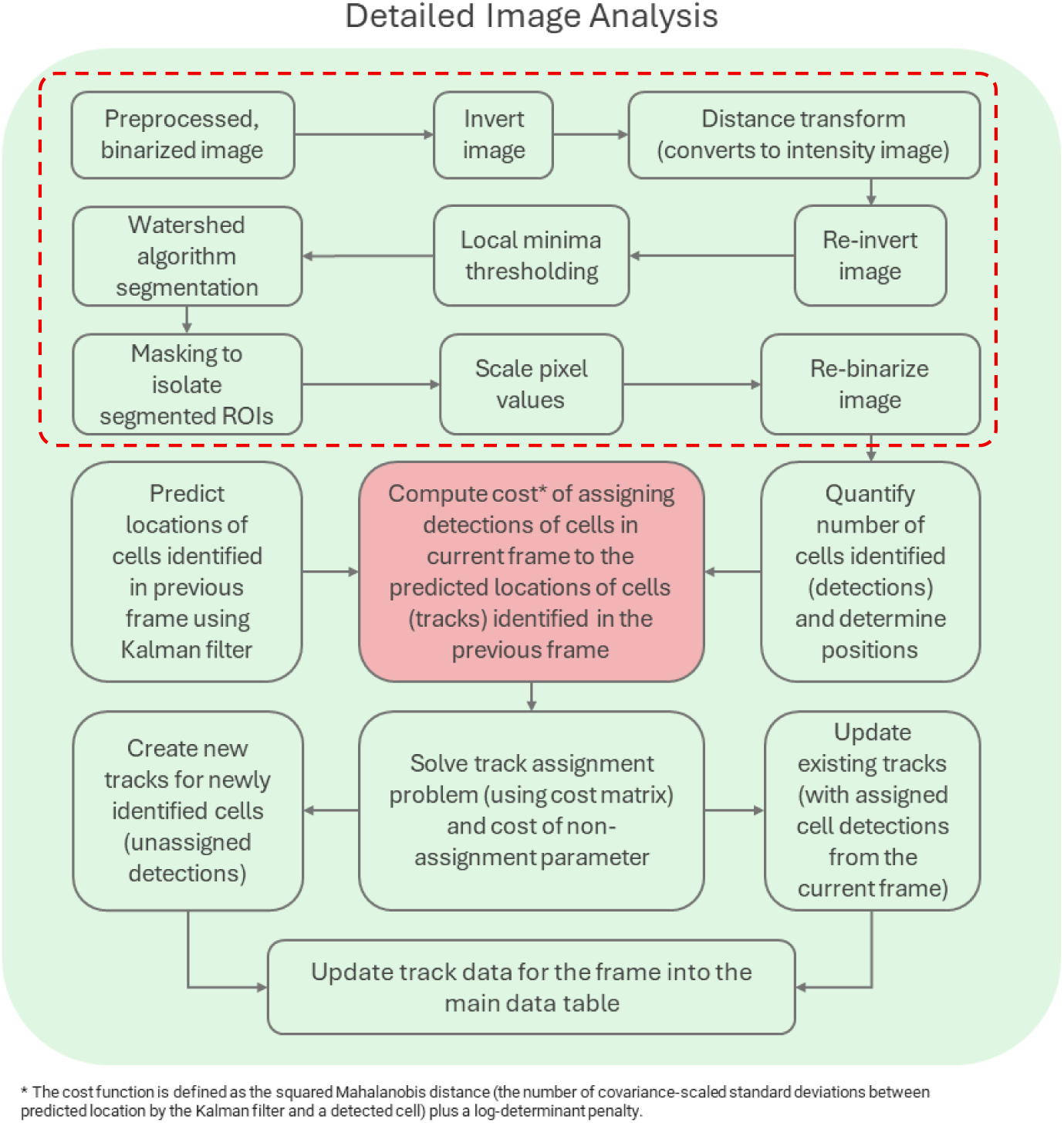
Image analysis pipeline for cell identification, segmentation, and quantification. Pre-processed fluorescence images were segmented using distance transformation and watershed-based object separation to identify individual cells. Cell positions were subsequently linked across sequential frames using a state-space tracking model comprising Kalman filter prediction, detection assignment, and track updating, enabling automated reconstruction of migration trajectories and downstream quantification of cell migration behaviour.

**Supplementary Figure S3.**
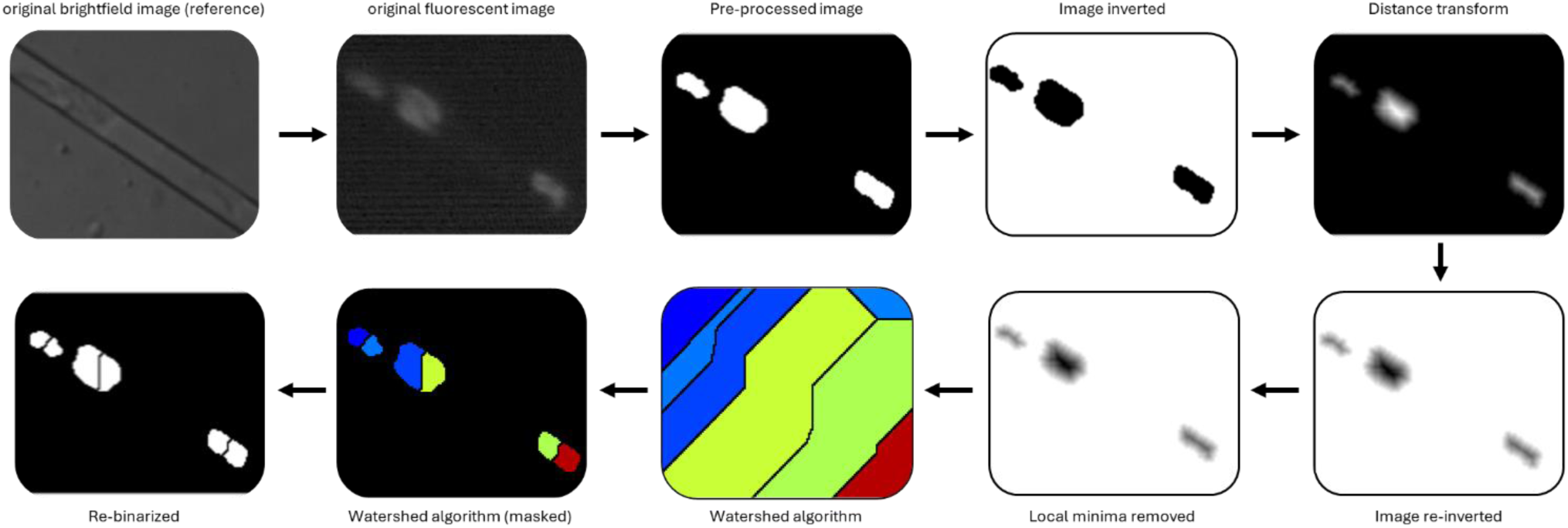
Watershed algorithm for distinguishing between adjacent cells. Original fluorescence images were pre-processed and inverted prior to application of a distance transform. The transformed image was subsequently re-inverted, local minima were removed to minimise over-segmentation, and a watershed algorithm was applied to separate neighbouring cells. The resulting segmentation mask was re-binarized to generate individual cell objects for downstream tracking and analysis.

**Supplementary Figure S4.**
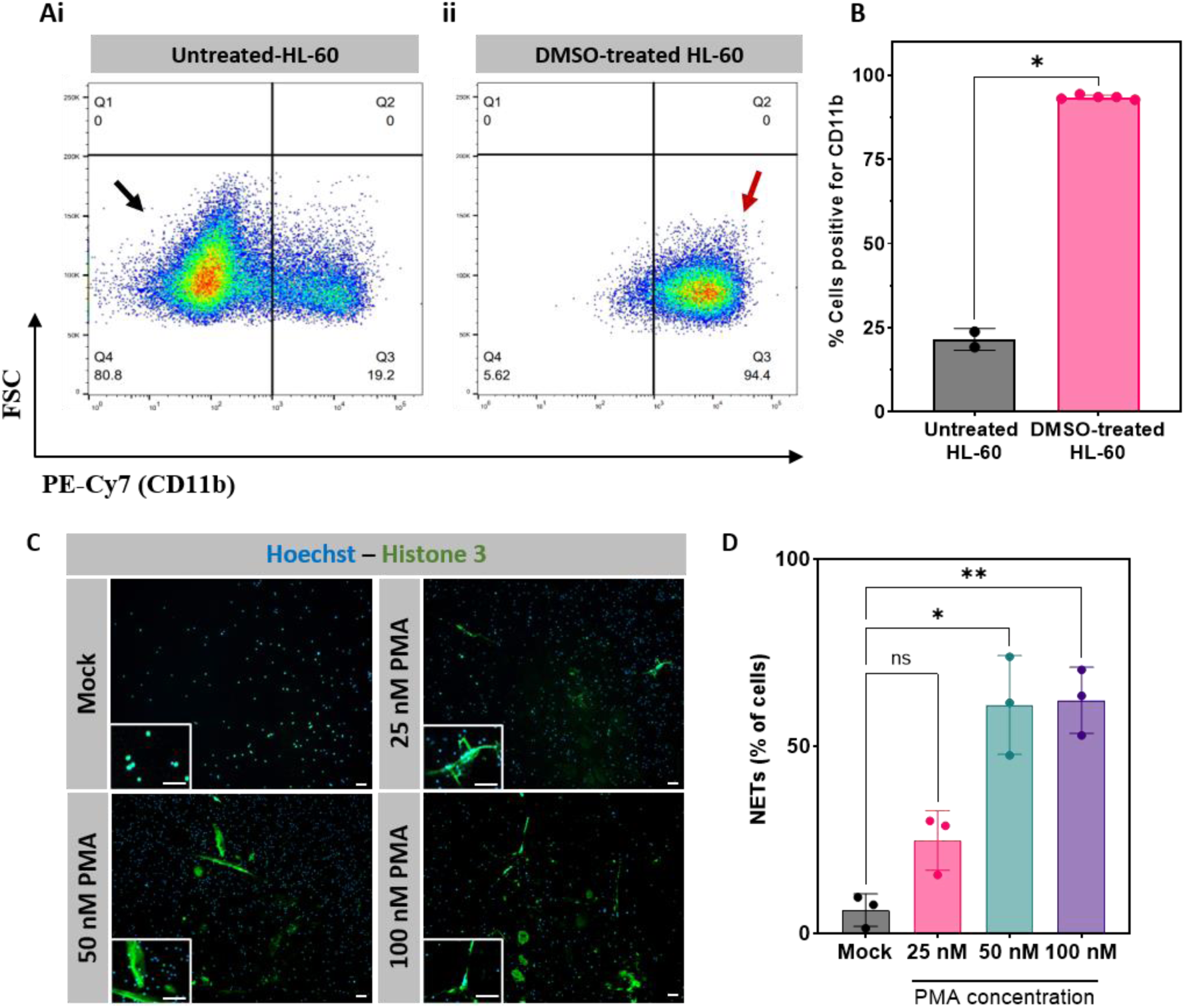
(A) Flow cytometry dot plots of HL-60 cells cultured without (i) or with (ii) differentiation media containing 1.3% DMSO. Arrows indicate CD11b-negative (black) and CD11b-positive (red) populations. (B) Summary of CD11b-positive cell populations. Statistical analysis performed using an unpaired t-test. *p < 0.05. (C) Representative fluorescence images showing Hoechst-stained nuclei and anti-Histone 3 staining of NLCs after stimulation with 0-100 nM Phorbol 12-myristate 13-acetate (PMA), indicative of neutrophil extracellular trap (NET) formation. Scale bar = 200 µm. n = 3 cultures imaged, with one representative culture shown. (D) Quantification of NET formation, expressed as percentage (%) of cells forming NETs. Statistical analysis between unstimulated NLCs and PMA-stimulated NLCs was performed using One-way ANOVA, ns: not significant, **p < 0.01, ****p < 0.0001.

**Figure S5.**
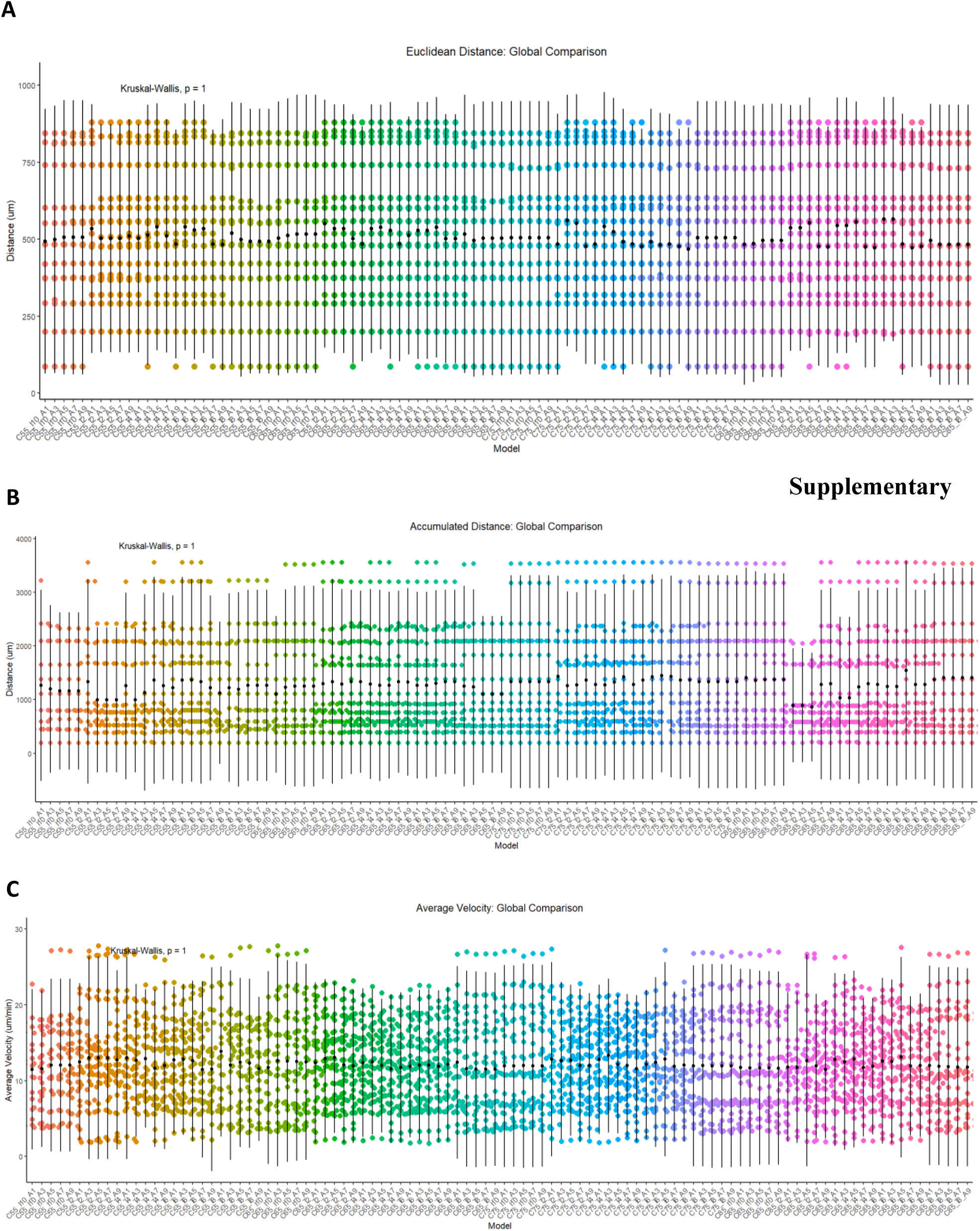
Key tracking parameters were systematically varied to generate 100 tracking models. End-point metrics from all models were compared across representative microchannels (channels 1, 5 and 7) over 180 imaging frames to identify parameter combinations that produced the most consistent trajectory reconstruction. (i) Final Euclidean distance, (ii) final accumulated distance, and (iii) final cell speed. Global comparisons across all 100 models were performed using the Kruskal–Wallis test, with no significant differences observed for any metric (all p = 1.0).

**Supplementary Figure S6.**
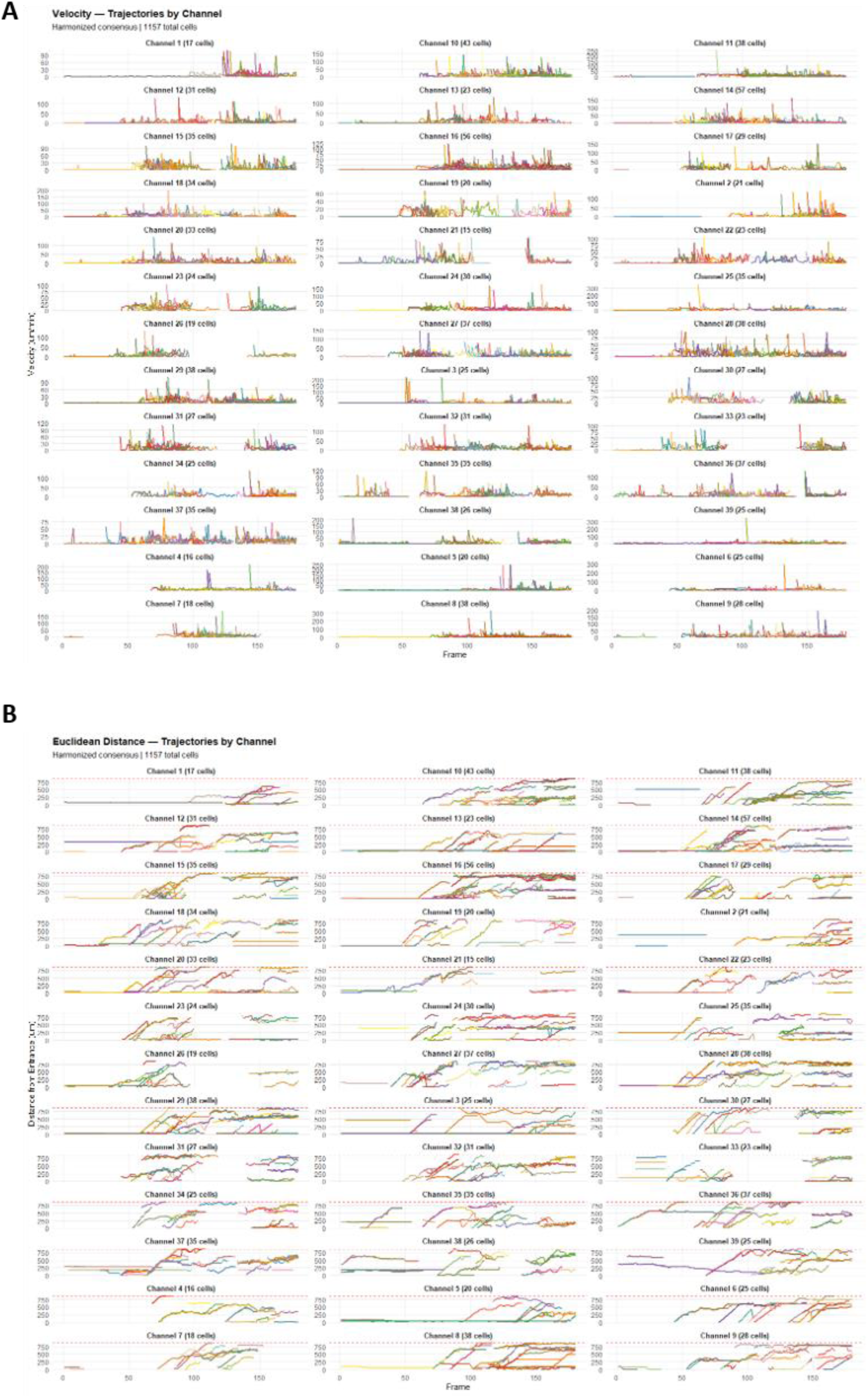
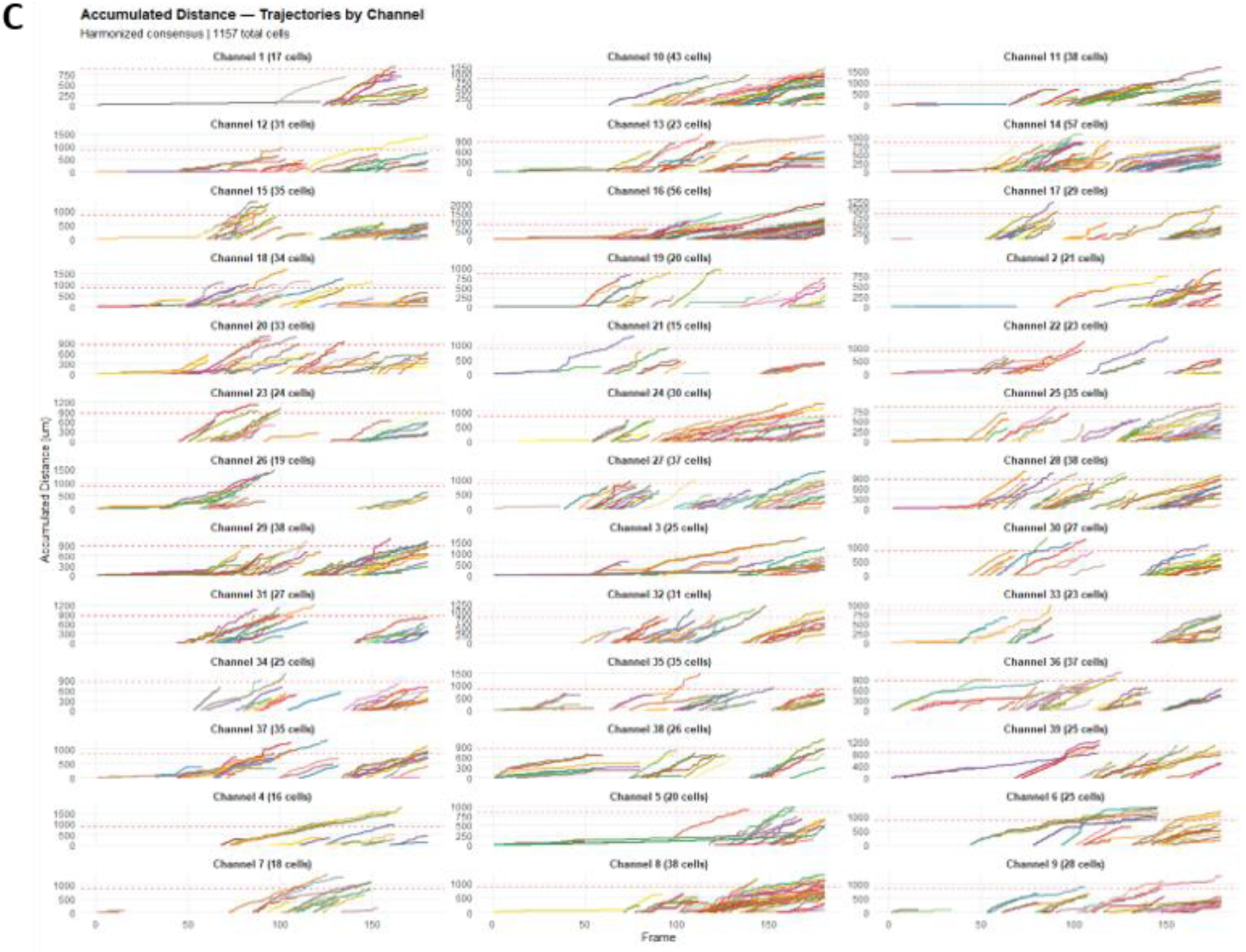
Individual cell migration metrics generated using the scAIR-ML platform from the fMLP dataset, including (A) velocity, (B) Euclidean distance, and (C) accumulated distance.

**Supplementary Figure S7.**
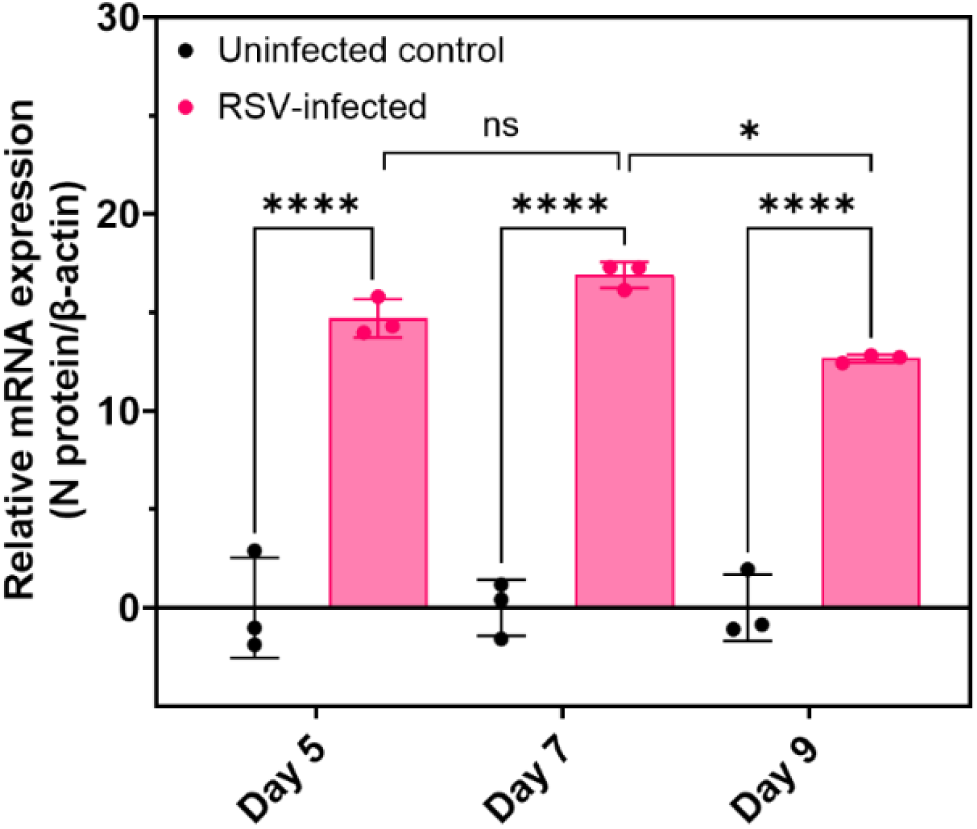
RSV infection of AECs in standard Transwell^®^ culture. mRNA expression of RSV N protein at days 5, 7, and 9 p.i. *n* = 3 cultures per timepoint. Statistical significance determined using Two-way ANOVA with multiple comparisons. *p < 0.05, ****p ≤ 0.0001.

**Supplementary Figure S8.**
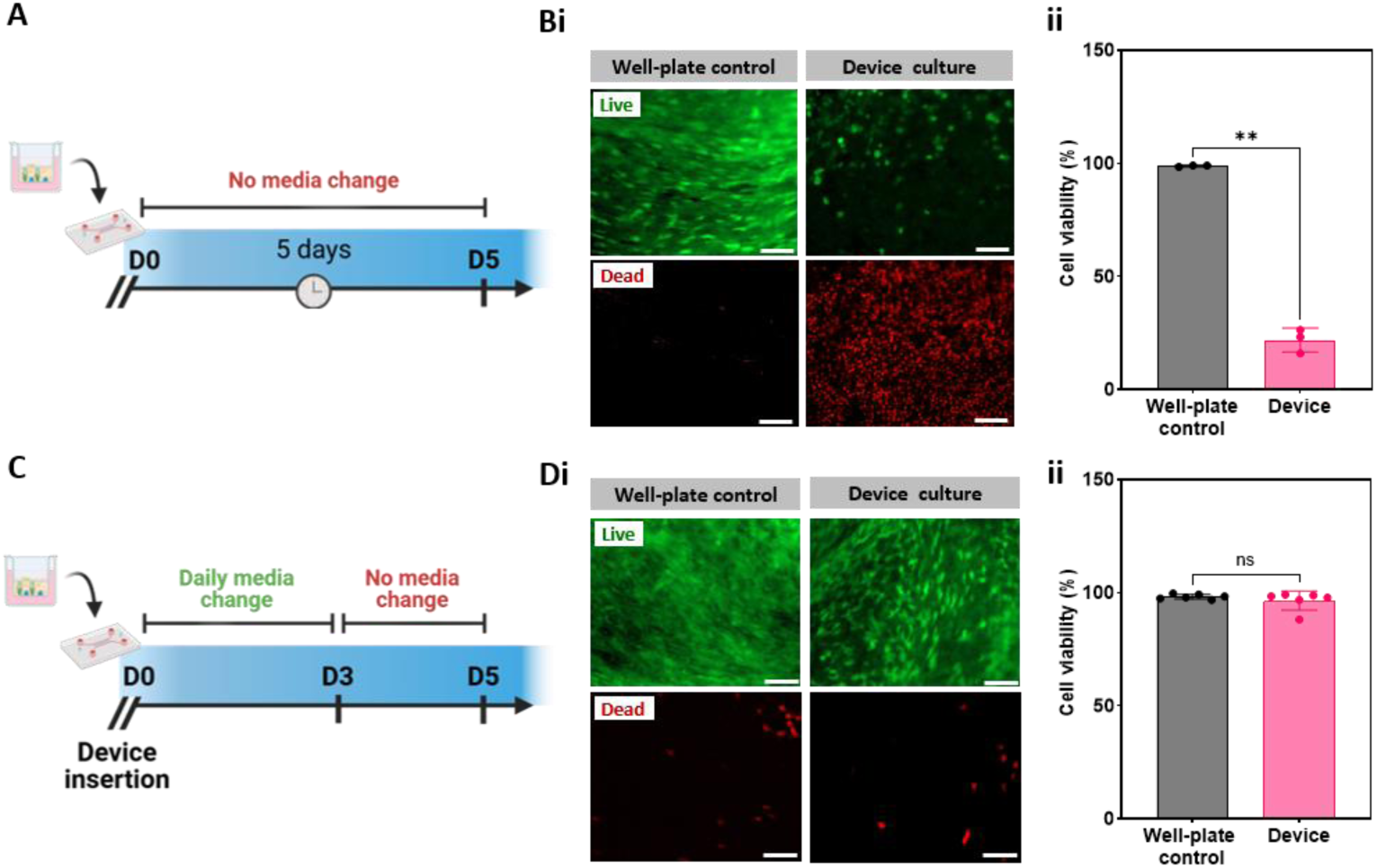
Comparison of AEC viability in scAIR devices cultured with different media replenishment protocols. (A) Experimental timeline of AECs transferred to devices on day 25 post-seed (day 0 in schematic), with no media replenishment for 5 days. (B)(i) Live (FDA)/dead (PI) staining of HBECs at day 5. (ii) Cell viability quantification as a percentage (%) of live cells (C) Experimental timeline of AECs transferred to devices on day 25 post-seed (day 0 in schematic), with media replenished every 24 hours for 3 days, followed by a 48-hour period without media change. (D)(i) Live (FDA)/dead (PI) staining of HBECs at day 5. (ii) Quantification of cell viability (% live cells). *n*=3 cultures per condition, with n=2 experimental replicates. Scale bar = 100 µm. Statistical significance determined using an unpaired t-test. ns: not significant, **p < 0.01.

**Supplementary Figure S9.**
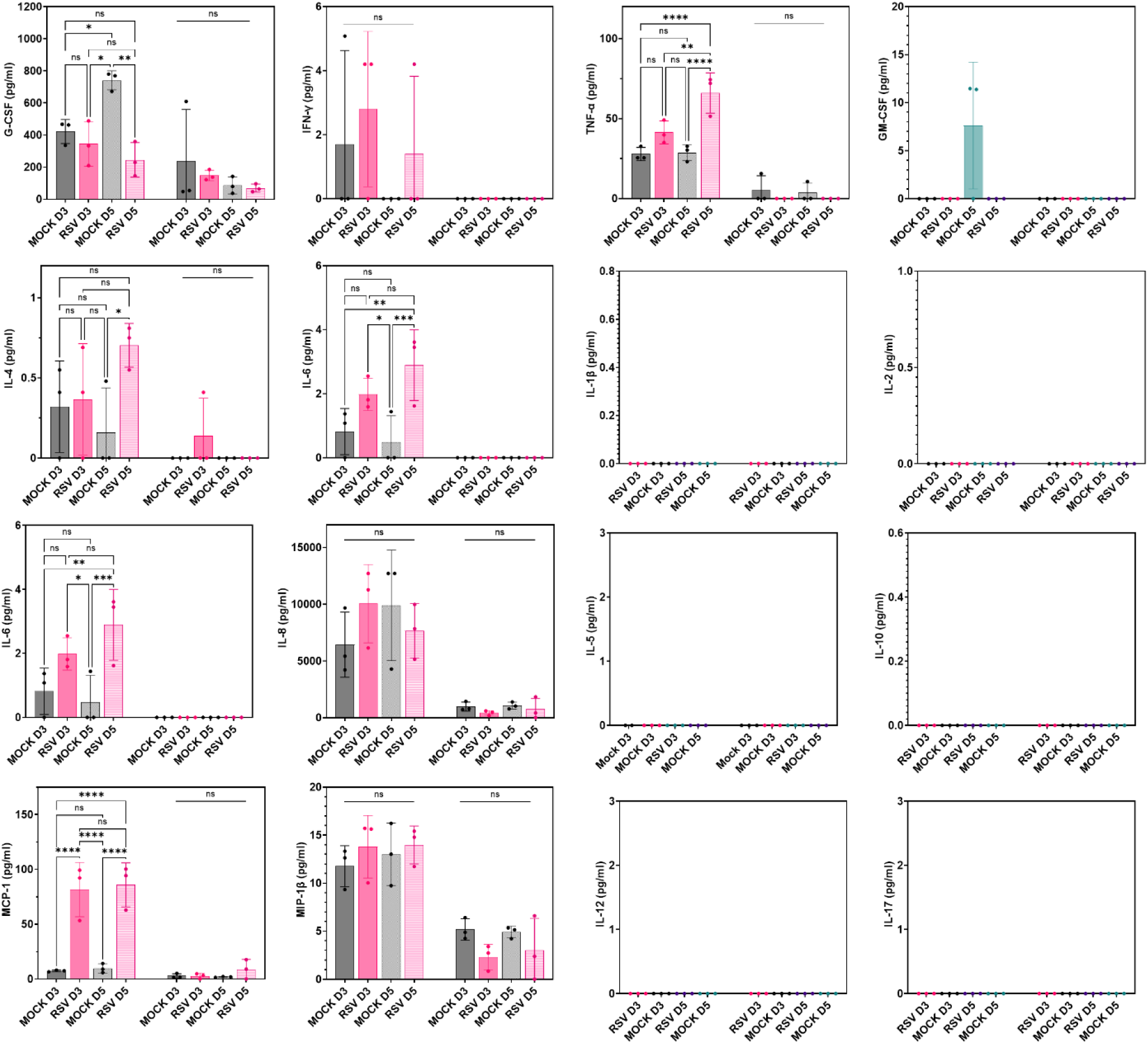
Cytokine and chemokine secretion profiles of basal media in scAIR devices following RSV infection of AECs at days 3 and 5 p.i. *n*=3 scAIR devices per condition. Statistical significance determined using Two-way ANOVA with multiple comparisons. ns: not significant, *p < 0.05, **p < 0.01, ***p < 0.001, ****p < 0.0001.

**Supplementary Figure S10.**
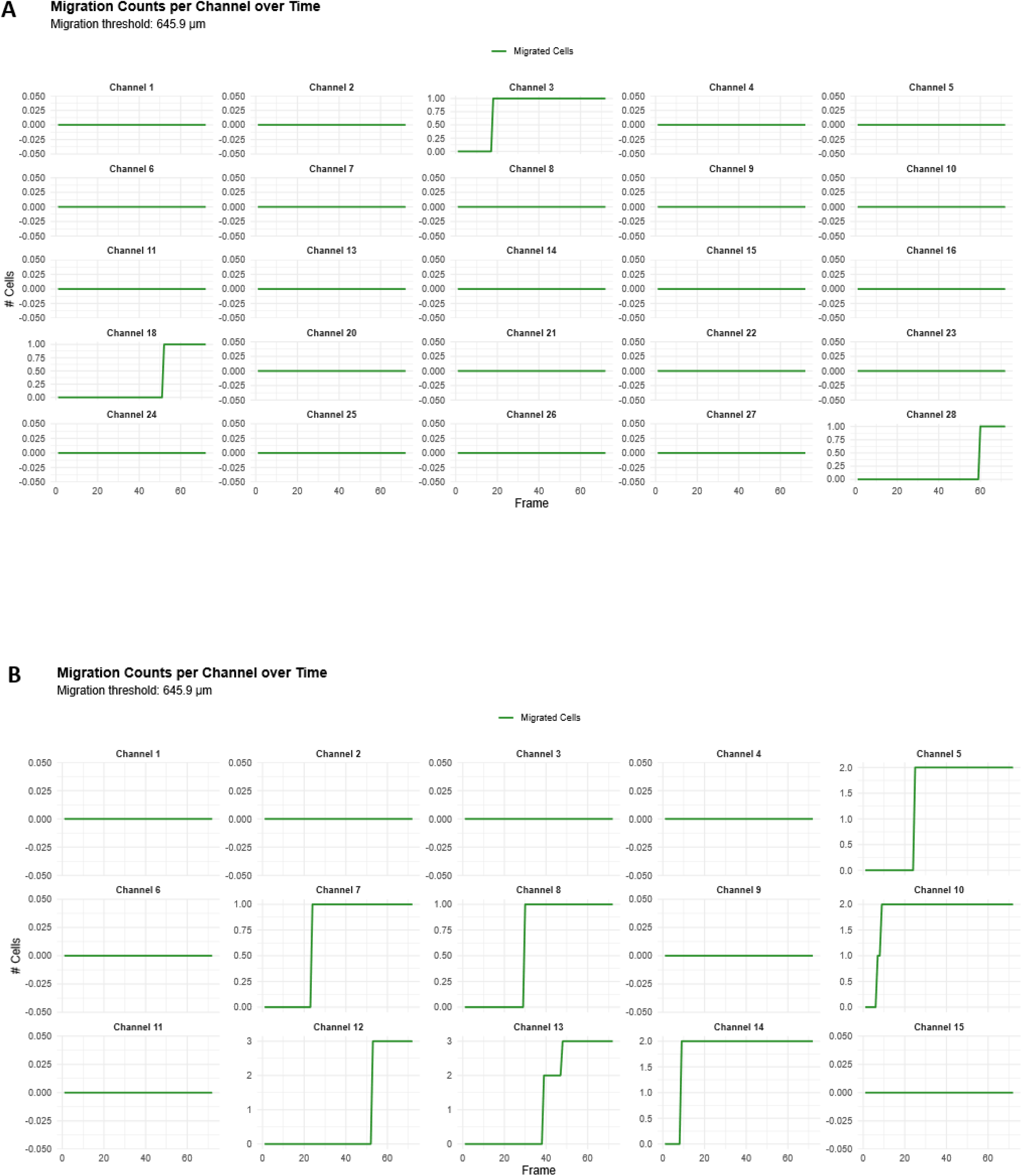
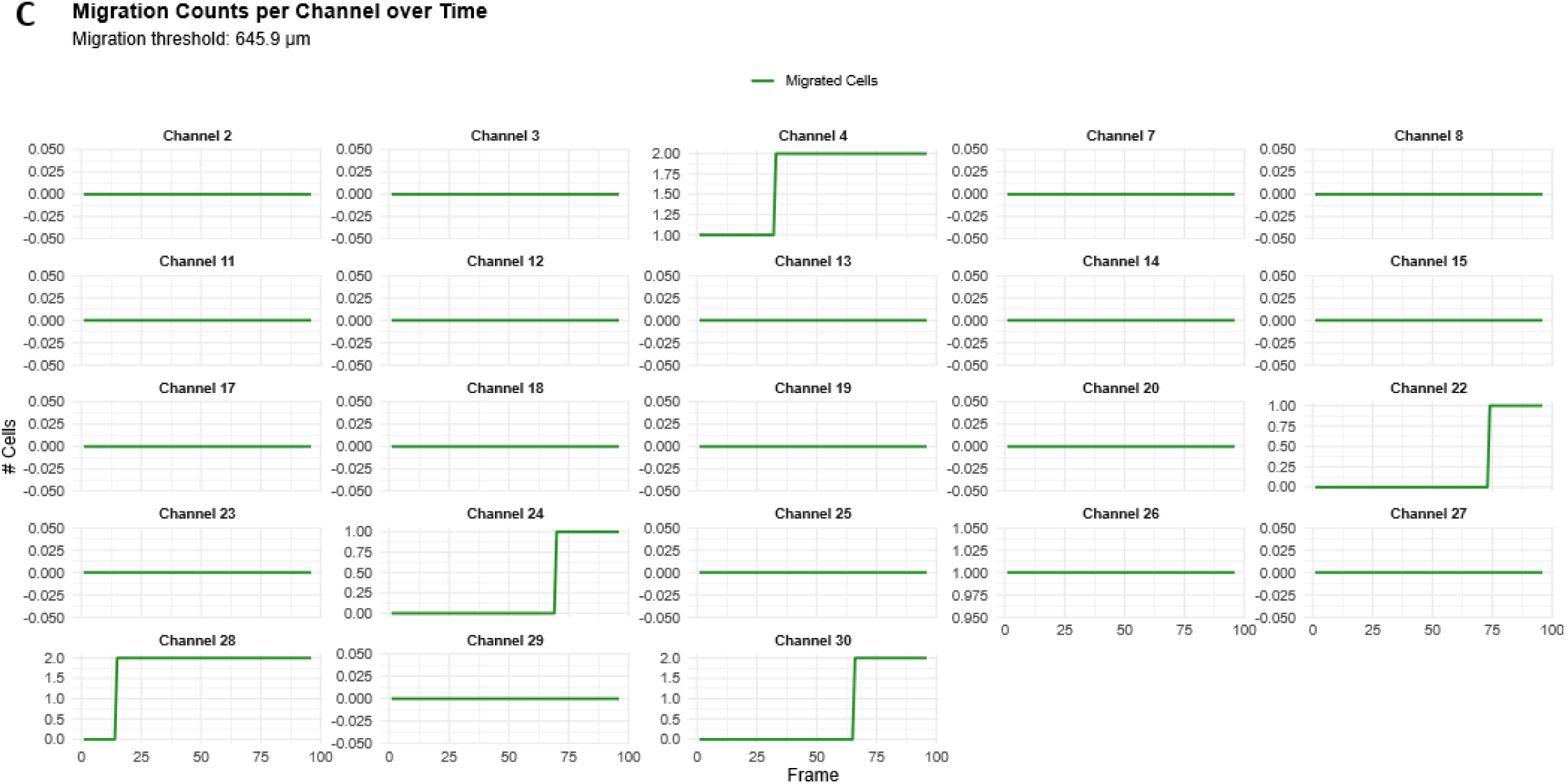
Cell migration metrics generated using the in-house MLA from (A) Uninfected, (B) +RSV, and (C) + RSV, +mAb datasets. Each miniature graph represents a single microchannel, with green lines representing accumulated migration metrics within that microchannel over time.

**Supplementary Table S1.** Settings used for flow cytometry experiments.

|  |  |
| --- | --- |
| Voltage settings | SSC: 451 |
|  | FSC: 319 |
|  | BV510: 341 |
|  | PE-Cy7: 462 |
| Loader settings | Sample Rate: 12 |
|  | Sample Volume: 300 |
| Event settings | Collect 50,000 total events per sample |

**Supplementary Table S2.** Raw live-cell imaging dataset information.

| Dataset | Image Resolution (pixels) [$\mu\text{m}$] | Bit-depth (bits) | Length of live-cell imaging experiment (hours) | Time interval (min) | Number of Frames | Number of Channels |
| --- | --- | --- | --- | --- | --- | --- |
| FMLP1 | $2630 \times 3886$<br>[$1675 \times 2475$] | 8 | 10 | 2 | 180 | 39 |
| RSV1 | 3854 × 1946<br>[2455 × 1240] | 8 | 6 | 5 | 72 | 15 |
| RSV2 | 3899 × 1967<br>[2484 × 1253] | 8 | 6 | 5 | 72 | 17 |
| MOCK1 | 3855 × 2882<br>[×] | 8 | 6 | 5 | 72 | 28 |
| MOCK2 | 3908 × 2969<br>[×] | 8 | 6 | 5 | 72 | 29 |
| Anti-TNFa1 | 2668 × 2896<br>[×] | 8 | 6 | 5 | 72 | 28 |
| Anti-TNFa2 | 2670 × 2910<br>[×] | 8 | 6 | 5 | 72 | 30 |

**Supplementary Table S3.**
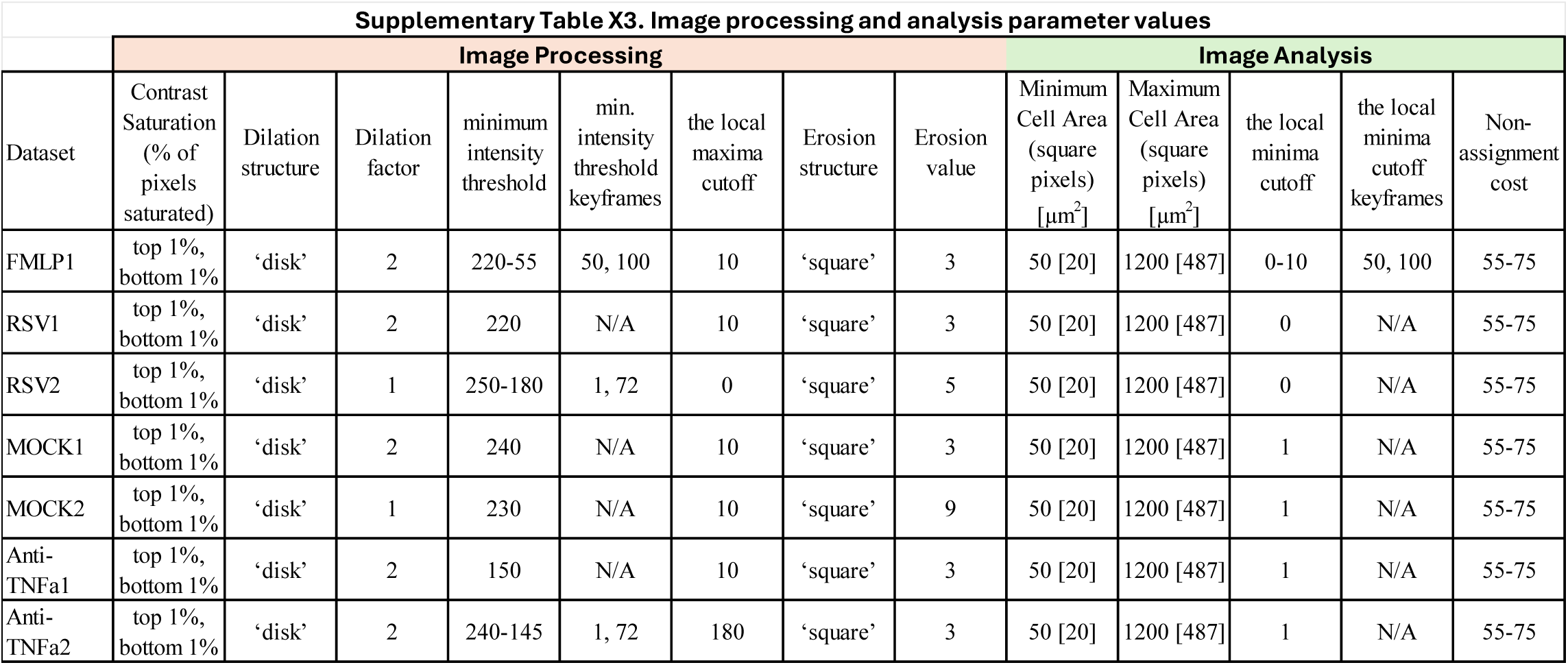
Machine learning image processing parameter values.

**Supplementary Table S4.** Classification system used to define behavioural cell movement patterns. Net progress, the net displacement of cells from their starting to final positions; tortuosity, the degree of deviation of cell trajectories from a straight path; and range explored, the spatial extent of the area explored by cells during the tracking period.

| Category | Net progress | Tortuosity | Range explored (μm) | Qualitative description |
| --- | --- | --- | --- | --- |
| Forward moving | high positive (≥100) | low (1-5) | ≥30 | strong movement towards exit |
| Backward moving | high negative (≤-50) | low (1-5) | ≥30 | strong movement away from exit |
| Oscillatory | low<br>($<100$) | High<br>($>5$) | $\geq 30$ | active motility with<br>little net progression |
| Low motility | near-zero<br>($<30$) | varies | $<30$ | minimal cell movement |
| Mixed | varies | varies | varies | motility profile fits<br>several different<br>categories |

**Supplementary Table S5.**
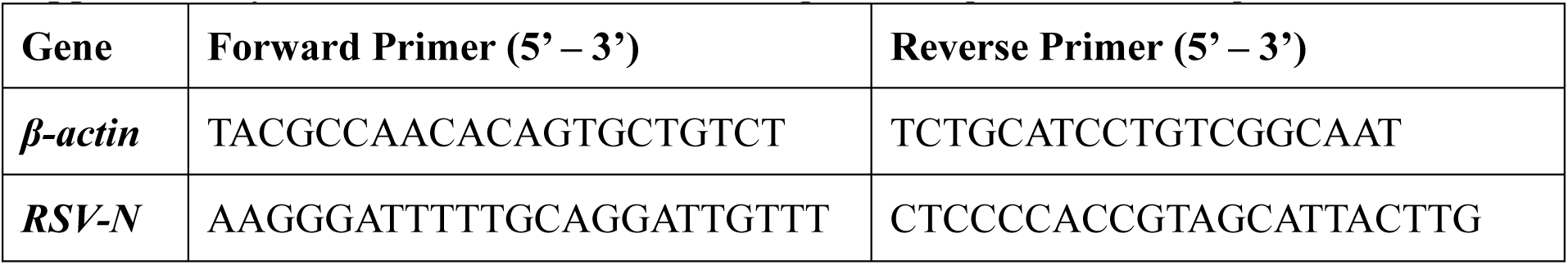
Forward and reverse primer sequences for RT-qPCR.

**Supplementary Movie S1**: NLC migration in response to exogenous fMLP. Codec: Motion JPEG Video (MJPG).

**Supplementary Movie S2 & S3**: NLC migration in response to uninfected AECs. Codec: Motion JPEG Video (MJPG)

**Supplementary Movie S4 & S5**: NLC migration in response to RSV-infected AECs. Codec: Motion JPEG Video (MJPG)

**Supplementary Movie S6 & S7**: NLC migration in response to RSV-infected AECs pre-treated with TNFα-neutralising monoclonal antibody Adalimumab. Codec: Motion JPEG Video (MJPG)

